# Extensive and differential platinum chemotherapy mutagenesis in children

**DOI:** 10.1101/2025.10.12.681852

**Authors:** Anna Wenger, Henry Lee-Six, Manas Dave, Mehdi Layeghifard, Andrew R.J. Lawson, Federico Abascal, Pantelis A. Nicola, Taryn D. Treger, Toochi Ogbonnah, Conor Parks, Thomas R.W. Oliver, Jonathan Kennedy, Angus Hodder, Nathaniel D. Anderson, Felipe Luz Torres Silva, Mi K. Trinh, Thomas Dowe, Marwo Habarwaa, James J. Sun, Sergio Assia-Zamora, Miriam Cortes-Cerisuelo, Wayel Jassem, Charlotte Town, Anil Dhawan, Vandana Jain, Karin Straathof, Maesha Deheragoda, Iñigo Martincorena, Liina Palm, J. Ciaran Hutchinson, Tim H.H. Coorens, Claire Trayers, Nigel Heaton, Adam Shlien, Yoh Zen, Foad J. Rouhani, Sam Behjati

**Affiliations:** Wellcome Sanger Institute; Hinxton, CB10 1SA, UK; Cambridge University Hospitals NHS Foundation Trust; Cambridge, CB2 0QQ, UK; Program in Genetics and Genome Biology, The Hospital for Sick Children; Toronto, ON, Canada; Department of Paediatrics, University of Cambridge; Cambridge, UK; Great Ormond Street Hospital for Children; London, United Kingdom; Postgraduate program in Genetics and Molecular Biology, Institute of Biology, State University of Campinas (UNICAMP); Campinas, São Paulo, Brazil; Liver Transplant Unit, King’s College Hospital NHS Foundation Trust; London, UK; The Francis Crick Institute; London, UK; King’s Health Partners Centre for Translational Medicine; London, UK; Pediatric Liver, GI and Nutrition Center and Mowat Labs, King’s College Hospital; London, UK; University College London (UCL) Cancer Institute, Great Ormond Street NIHR Biomedical Research Centre; London, UK; Broad Institute of MIT and Harvard; Cambridge, 02142 MA, USA; Laboratory Medicine and Pathobiology, University of Toronto; Toronto, ON, Canada; The Roger Williams Institute of Liver Studies, King’s College London; London, UK

## Abstract

Childhood cancer survivors often develop long-term adverse effects, which may be caused by direct mutagenesis of cytotoxic agents. Some of these agents generate distinctive DNA imprints (mutational signatures), as exemplified by platinum chemotherapeutics. Here, we examined chemotherapy mutagenesis in paediatric tissues by deploying a duplex sequencing method (NanoSeq), which enables mutation calling from single DNA molecules. We surveyed whole genomes of paediatric liver, blood and other tissues, obtained from surgical resections and at post-mortem. Platinum signatures pervaded all tissues extensively, elevating mutation burdens of paediatric tissues to levels seen in adults. Remarkably, we found a tissue-specific mutational signature in the liver. We examined the functional potential of mutations by gene focused NanoSeq, which revealed that platinum agents cause a vast repertoire of cancer causing variants across normal tissues, such as leukaemogenic mutations in blood. This finding may conceivably link cancer treatment in childhood to mutation-driven long term sequelae.

## INTRODUCTION

Some chemotherapeutics confer cytotoxic effects through damage of DNA^1^. When this damage is not lethal to cells, agent-specific mutational imprints (signatures)^2^ may emerge in surviving cells if DNA repair is imperfect. Investigations of the past decade have documented a variety of chemotherapy mutational signatures in both cancer (typically in the form of recurrences or metastases) and clonal expansions of normal cells^2–7^. Chemotherapy mutagenesis in normal tissues has mainly been studied in blood through high depth sequencing of haematopoietic cancer genes. This approach has identified clonal expansions (clonal haematopoiesis)^8^ which, in some adult individuals with prior cancer treatment, were characterised by mutational signatures attributed to specific chemotherapy agents^9,10^.Chemotherapy mutagenesis has been further substantiated by DNA sequencing of individual blood cells, expanded through single cell cloning for high fidelity DNA amplification, demonstrating that it pervades blood cells even in the absence of clonal expansions^11,12^. Similarly, investigations of genomes of cancer relapses, which are reporters of past treatment and exposures (**Fig. 1A**), have corroborated mutational signatures attributable to chemotherapy, in particular to platinum-based agents^3,4^. Recent studies of tissues other than blood indicate that chemotherapy mutagenesis affects human tissues more broadly^6,13–16^. It is unknown whether the same exposure generates different mutations across tissues within an individual. This is a particularly pertinent question in children in whom the consequences of potentially harmful mutagenesis, such as second cancers, may unfold over the span of a lifetime^17–19^.

**Fig. 1.**
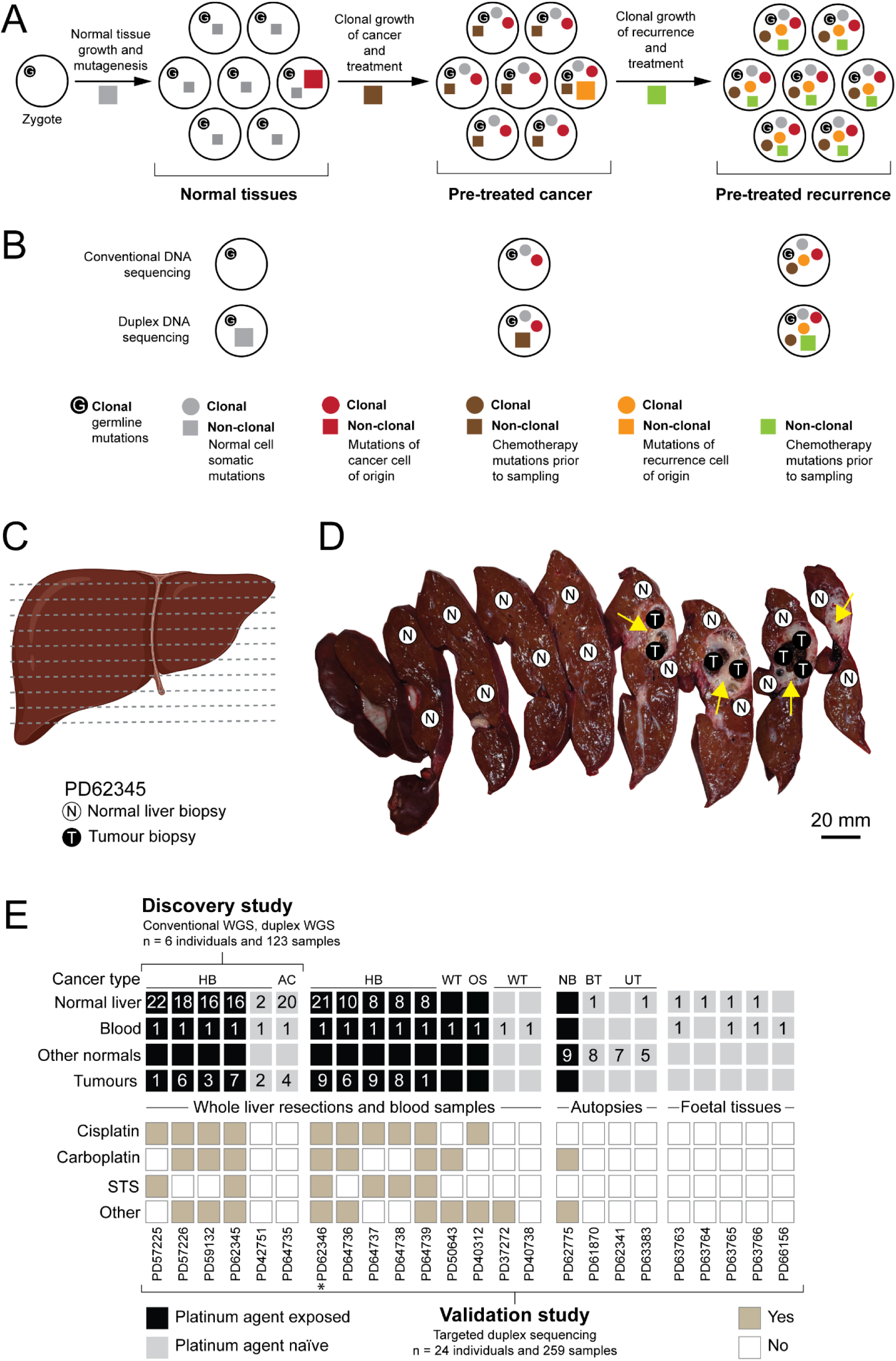
Conventional vs duplex sequencing, and study overview. **(A)** Schematic showing how mutations are accrued from the fertilised egg to a tumour recurrence. The fertilised egg harbours germline mutations which pervade all cells of the body. Normal cells acquire somatic mutations as they divide due to imperfect DNA replication (grey squares). Cancer causing mutations (red square) may transform one of these cells, which then becomes the cancer cell of origin that clonally expands into a tumour, and the somatically acquired mutations thus become clonal (grey dot). All tumour cells carry the mutations of the cancer cell of origin (red dot) and acquire additional cell-specific (non-clonal) mutations as they diversify and from mutagenic chemotherapy (brown square). One of the cells will be the recurrence cell of origin and grow clonally into the recurrent tumour and all cells will therefore carry the mutations from that cell (orange dot), and will gain private non-clonal mutations induced by the additional treatment (green square). **(B)** The mutations that are detected by conventional DNA sequencing (top) and duplex DNA sequencing (bottom) in normal tissues, pre-treated cancer and pre-treated recurrence respectively. Note that non-clonal mutations (present in individual cells) can be detected by duplex sequencing whereas mutations need to become clonal to be detected by conventional DNA sequencing (e.g. the non-clonal somatic mutations, grey square, present in the cancer cell of origin will become clonal in the pre-treated cancer, grey dot, and thus detectable by conventional DNA sequencing). **(C)** Liver schematic of sampling of PD62345. Created with Biorender. **(D)** Picture shows the fresh, unfixed liver of PD62345 after resection and prior to sampling. The sampling strategy is shown with each circle representing the site of tissue sampling. Normal liver (N) and tumour (T) tissue. Yellow arrows delineate the tumour. **(E)** Overview of samples and children in the study detailing the number and type of samples, sequencing assays and the treatment for each child. The blood sample for PD57226 was not taken at the same time point as the liver was sampled. HB: hepatoblastoma, AC: metastatic adrenocortical carcinoma in the liver, WT: Wilms tumour, OS: osteosarcoma, NB: neuroblastoma, BT: benign foetal lung tumour, UT: undifferentiated tumour, STS: sodium thiosulfate, Other: other treatment than cisplatin/carboplatin/STS. * PD62346 is a recurrent hepatoblastoma.

The platinum-based agents cisplatin and carboplatin have emerged as principal exemplars of mutation-inducing chemotherapeutics^4,20^. They are the backbone of many treatment protocols, particularly of solid childhood tumours. Unlike with other mutagenic agents (such as the purine analogue, thiopurine), mutagenesis by cisplatin and carboplatin is independent of cell cycle^21–23^ and seems to be pervasive in both adult normal and cancerous tissues following treatment^2,3,6^. The extent and nature of platinum agent mutagenesis in children remain largely unknown. The largest study to date, examining 2860 childhood cancer survivors, found no platinum signals in blood^24^ whilst another study of cancer survivors, which included some children, found only subtle effects^25^. By contrast, there have been occasional reports of platinum agent mutagenesis in blood^23,26^ and other normal tissues^13^ of exposed children. Furthermore, studies of recurrent or second cancers in children have demonstrated mutagenesis by cisplatin and carboplatin^13,26–29^. The discrepancy can probably be attributed to technological limitations of conventional targeted DNA sequencing, which depends on the presence of clonal outgrowths for mutation detection (**Fig. 1A-B**)^30^. This shortcoming can be overcome by methods that enable precise mutation calling from single DNA molecules^31^, independent of clonal expansions, through barcoding of both strands of DNA (duplex sequencing) to differentiate true mutations (present on both strands) from artefacts (present on one strand) (**Fig. 1A-B**). Using this approach, we examined chemotherapy mutagenesis in children, focusing on liver and blood, supplemented with a variety of other normal tissues obtained at post mortem and surgical resections.

## RESULTS

### Overview of experiment

Initially, our investigation focused on children with hepatoblastoma who underwent whole liver replacements as part of their treatment, which enabled us to study from the same child multiple areas of normal liver, tumour, and matched blood (**Fig. 1C-D**). Prior to surgery, children with hepatoblastoma are typically treated with cisplatin, either alone or in combination with carboplatin and other agents. Our study also included children who had received non-platinum agents as well as treatment-naïve children, whose liver tissues we sampled from surgical specimens or at post-mortem. In addition, we studied human foetal livers as an additional treatment-naïve control (**Fig. 1E**). As these contain a large amount of hematopoietic islands, we were able to examine both liver and hematopoietic cells by separating respective cell populations from each other (see **Methods**). Our principal tool for mutation detection was an established implementation of duplex sequencing (NanoSeq)^31^, to examine genome wide mutations (whole genome NanoSeq), as well as mutations within specific genes (targeted NanoSeq^32^). The latter enabled us to determine potential functional consequences of variants. We called mutations using established variant calling pipelines^31–33^ (detailed in **Methods**). An overview of the tissues we studied and our experimental strategy is shown in **Fig. 1E** and is detailed in **table S1.**

### Increased mutation burden in exposed liver tissues

In a first experiment we studied normal livers (n=93 samples), blood (n=6 samples) and corresponding tumours (19 hepatoblastoma samples and 4 liver metastasis samples of an adrenocortical carcinoma) of 4 pre-treated and 2 chemotherapy-naïve children. Within the normal liver we systematically sampled all anatomical regions (**Fig. 1C-D**) and interrogated DNA by conventional whole genome sequencing (WGS) and by whole genome NanoSeq. Conventional WGS of normal liver and blood yielded, as expected, occasional base substitutions consistent with mosaic variants, with an average of 14 substitutions (range 2-31) per sample (**Fig. 2A; table S2**). Conventional WGS of the tumour showed typical cancer genomes with hundreds of substitutions as well as other expected somatic changes (insertions/deletions (indels), common copy-number changes for hepatoblastoma such as chromosome 11p LOH) including typical cancer-causing (driver) mutations (**fig. S1**; **table S3**). By contrast, whole genome NanoSeq revealed an extraordinarily high substitution burden across all normal liver tissues, blood, and tumours, as well as additional variants in tumours (**Fig. 2B; table S4**). Compared to mutation burdens of corresponding untreated tissues, there was a several fold increase in substitution load in chemotherapy-treated tissues (9-fold for blood, 12-fold for normal liver, 5-fold for liver tumours), elevating the load of platinum-exposed tissues to that of adult liver^34^ and blood^35^ (**Fig. 2B**). Similarly, we observed increased burdens of indels and double base substitutions (DBS) in exposed normal tissues and tumours (**fig. S2A-C**). There were no systematic regional differences in substitution load within the normal liver except for in one case (PD57226, **fig. S2D-E**).

**Fig. 2.**
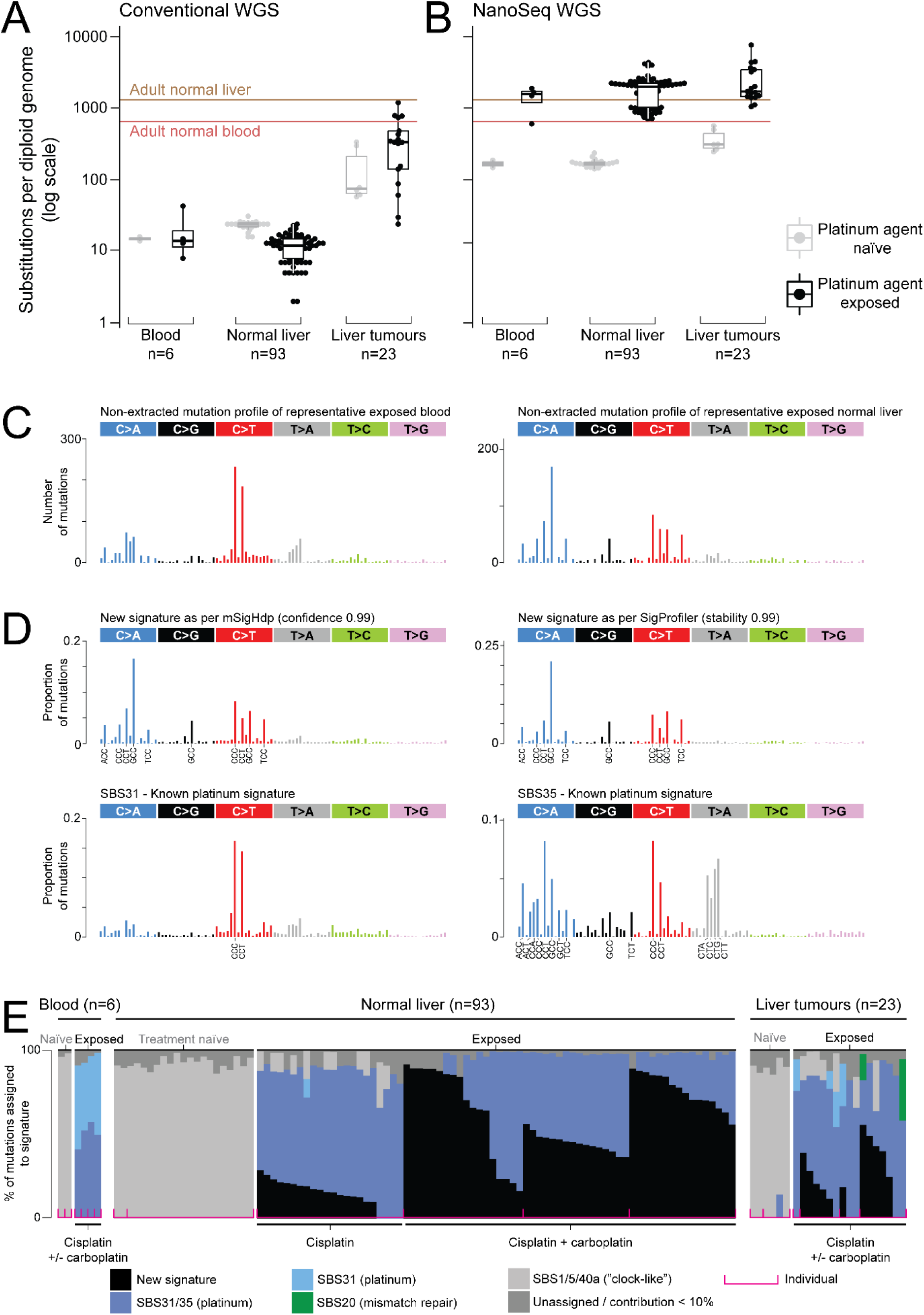
Whole genome mutation burden and signature analysis in discovery cohort. **(A)** Mutation burden by conventional whole genome sequencing (WGS). X-axis: different tissues. Y-axis: number of base substitutions per diploid genome (log scale). Red and brown horizontal lines represent mutation burdens observed in normal adult blood^35^ and liver^34^. **(B)** Mutation burden by WGS NanoSeq. Axes and horizontal lines as per (A). **(C)** Plots showing the normalised trinucleotide context (i.e. the reference base before and after a variant) of each base substitution (X-axis) and the frequency of each trinucleotide for platinum-exposed blood (left), and platinum-exposed normal liver sample (right) differ visually, and according to the *de novo* signature extractions (see (D)-(E)) where the new signature was absent from the blood sample while up to 92% of the mutations per normal liver sample assigned to the new signature. **(D)** Trinucleotide context for: Upper plots – a new signature as extracted from NanoSeq WGS data by mSigHdp (left) and SigProfilerExtractor (right). Lower plots - Known signatures of platinum agents (SBS31 and SBS35). **(E)** Contribution of signatures to each sample. X-axis: Each stacked bar plot represents a tissue sample as per labelling. Y-axis: % of mutations assigned to signature. Pink square brackets delineate individuals. Colours represent mutational signatures as per legend. The mSigHdp extraction is shown here. Similar results were obtained by SigProfilerExtractor (fig. S4B). Note that the new signature was only evident in platinum-exposed livers whereas conventional platinum signatures were pervasive across all platinum-exposed tissues. See representative spectra in (C).

### Distinctive mutagenesis in the liver

Next, we examined mutational signatures of single base substitutions (SBS), as defined by the trinucleotide context of substitutions, using two orthogonal analytical methods, mSigHdp^36^ and SigProfilerExtractor^37^. Conventional bulk WGS of tumour samples describes the history of the tumour (i.e. the most common ancestor cell the tumour bulk is derived from), but not the tumour’s current per cell mutational state (in the absence of treatment-related clonal expansions) (**Fig. 1A-B**). Accordingly, in mutation catalogues derived by conventional WGS, we did not find treatment-related signatures in pre-treated tumour samples. Instead, they exhibited mutation signatures SBS1 and SBS5, which are ubiquitous and have been attributed to endogenous processes^7^ (**fig. S3**). We could not extract signatures from normal blood and liver, due to the paucity of variants that conventional WGS delivered. In contrast, whole genome NanoSeq, which has single DNA molecule resolution and is independent of clones, revealed platinum agent mutagenesis (SBS31, SBS35) across pre-treated tissues, both normal and cancerous, which was even evident in non-extracted mutation profiles (**Fig. 2C-E**). In addition, in exposed normal liver and hepatoblastoma tumours, but not in blood, both methods extracted a new mutational signature (**Fig. 2D-E**; **table S5**), i.e. an imprint that could not be satisfactorily explained by the existing compendium of 86 signatures in COSMIC (**fig. S4A-B**) or by other platinum agent signatures documented in the literature (**fig. S4C**). The new signature was visible “by eye” in normalised trinucleotide spectra (i.e. non-extracted; **Fig. 2C**). It accounted for up to 92% of single base substitutions (range 0-92%; median 46%) in each exposed normal liver sample (**Fig. 2E**). Like other mutagen associated imprints, the new signature exhibited transcriptional strand bias (**fig. S4D**). Its characteristic features were C>A (blue), C>G (black) and C>T (red) mutations in a G<u>C</u>C trinucleotide context (**Fig. 2D**). The reliability of the extracted signature was high: 99% by both mSigHdp and SigProfilerExtractor. The signature analysis was otherwise unremarkable, bar several biopsies from one tumour (PD62345), which exhibited a signature (SBS20) proposed to be caused by defective mismatch repair. Signatures of indels (**fig. S5**) and DBS (**fig. S6**) showed patterns consistent with previous studies (*5*) of pre-treated (including platinum agents) tissues.

### Validation in 251 tissue samples by targeted NanoSeq

To validate and extend our findings, we deployed targeted NanoSeq, which enabled us to assess both mutation burden and the potential functional impact of variants in genes of interest. We targeted a total of 285 genes (**table S6**) comprising mostly cancer genes^38^ as well as 7 genes reported to harbour somatic mutations in metabolic liver disease of adults^39^. We first established whether targeted NanoSeq faithfully recapitulated findings from whole genome NanoSeq. We subjected the same stock DNA we had used for whole genome NanoSeq to targeted NanoSeq and found a high correlation of the mutation load (Pearson correlation=0.95, p=2.2×10^-16^; **fig. S7**).

We then applied targeted NanoSeq to a total of 251 tissue samples obtained from liver resections, paediatric post-mortems, and human foetuses, examining in total ∼108,000 diploid cells (251 tissues x median of 863 duplex coverage), and identified a total of 33,324 variants which we list in **table S7**. Other than blood and liver, we studied the following normal tissues: spleen, pancreas, thyroid, skin, adrenals, and heart. We performed signature assignment of the targeted NanoSeq data using two orthogonal methods, SigProfilerAssignment^40^ and mSigAct^41^, testing for the presence of the new signature, platinum signatures (SBS31, SBS35, SBS-carboplatin^5^) and “clock-like”^3^ signatures (SBS1, SBS5, SBS40a). Several patterns emerged from our analysis of mutation burden and corresponding signatures. First, platinum agent exposure increased the mutation load of all samples (**Fig. 3A**). Even short treatment durations increased the mutation load, as exemplified by an infant who received only a single cycle of carboplatin treatment (PD62775) which led to a 4.3 increase in single base substitutions in normal tissues compared to three platinum-naïve infants of a similar age (**Fig. 3B**). Second, chemotherapy mutation load increased with exposure, as, for example, evident in children who received cisplatin alone versus cisplatin and carboplatin combined (**Fig. 3A**). Third, in this validation cohort we found the new signature again evident in platinum-exposed liver (**Fig. 3C**), but not in exposed blood or in other paediatric tissues (adrenal, kidney, pancreas, spleen, thyroid, heart, skin, platinum-naïve liver). These included tissues exposed to carboplatin alone (PD62775) where signature analysis initially yielded conflicting results, likely resulting from low mutation counts detected by targeted NanoSeq. We resolved these discrepant findings through whole genome NanoSeq to obtain a greater number of variants (**Fig. 3D**). Signature analysis in this larger number of genome-wide mutations revealed a previously described carboplatin specific signature (SBS-carboplatin^5^) with no evidence of the new signature in these tissues (note that no liver sample was available from this child). Importantly, we also detected the new signature in tissues of children who had not received the chelating agent sodium thiosulfate, which is usually co-administered with cisplatin to protect hearing^42^, thus ruling out sodium thiosulfate as the cause of the new signature.

**Fig. 3.**
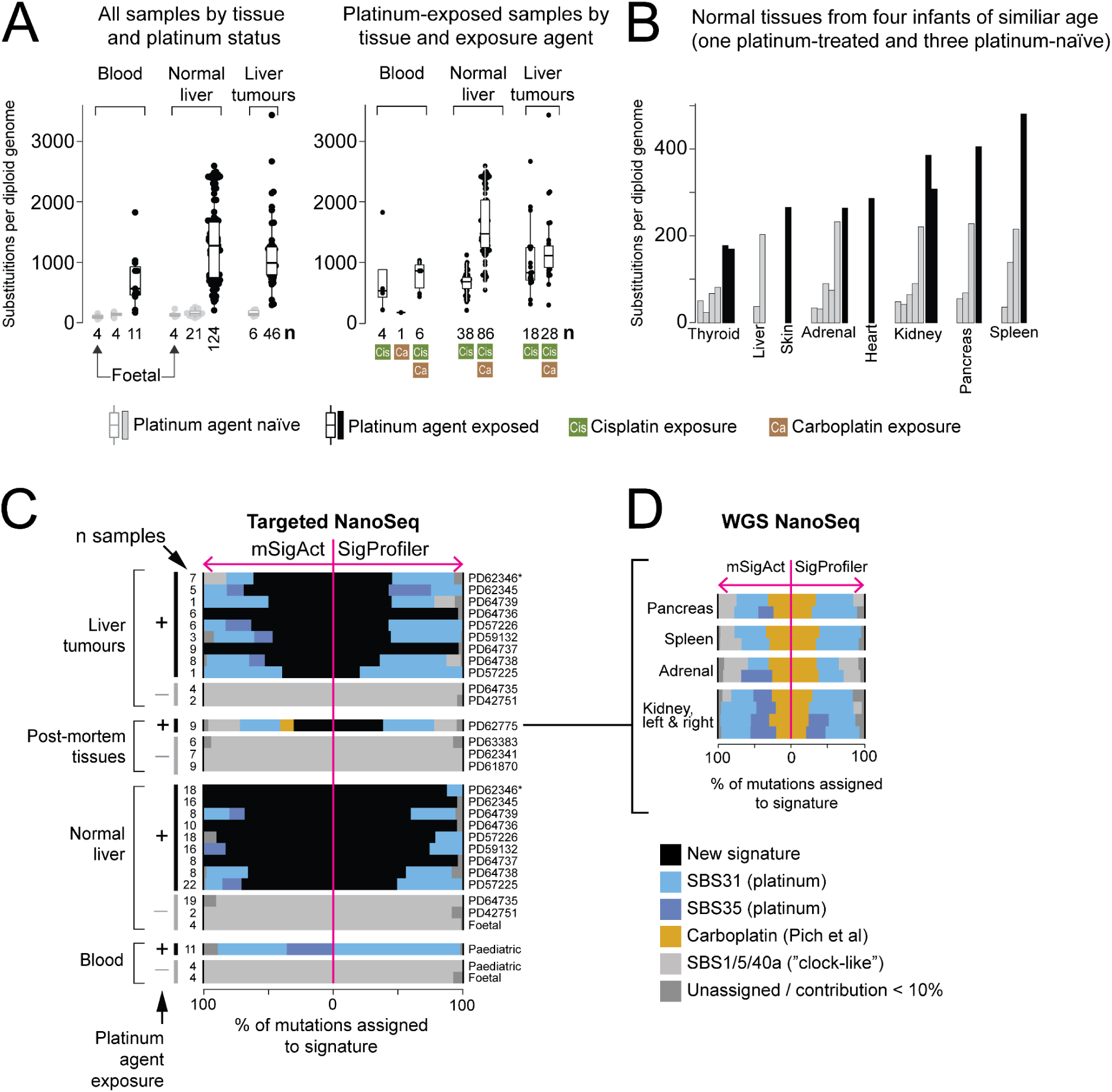
Mutation burden and signatures in validation cohort by targeted NanoSeq. **(A)** Targeted NanoSeq of 220 samples (including 120 samples from the discovery cohort). X-axis: Different tissues grouped as per labels in Figures. n=Numbers: number of samples per category. Y-axis: Number of single base substitutions per diploid genome. Exposure represented in colours (grey – platinum agent naïve; black – platinum agent exposed). Left: All 220 samples by tissue type. Right: 181 samples by platinum exposure. **(B)** Number of single base substitutions per diploid genome (Y-axis) in 31 different normal tissue samples obtained at post-mortems from four infants. The infants were of a similar age. Exposed tissues were obtained from one infant who had been treated with one cycle of carboplatin. Exposure represented in colours (grey – platinum agent naïve; black – platinum agent exposed). **(C)** Contribution of signatures to each sample. X-axis: % of mutations assigned to signatures with each signature represented by a different colour. Stacked bar plots to the left of the pink line are results from mSigAct whereas stacked bar plots to the right are results from SigProfilerAssignment. Y-axis: Each stacked bar plot represents a tissue sample as per labelling. Note that liver tissue was not available from PD62775. *PD62346 is a recurrent hepatoblastoma. **(D)** Re-examination of normal tissues from PD62775 by whole genome NanoSeq. Axes as per (C). Signature analysis of a greater number of NanoSeq WGS mutations resolves conflicting results of targeted NanoSeq (average of 58 mutations per sample by targeted NanoSeq vs average of 398 mutations per sample by NanoSeq WGS) and indicates that the new signature may be specific to platinum-exposed liver.

### Validation in external data sets

To further validate our finding of a new mutational signature, we examined external substitution catalogues derived from normal and cirrhotic adult liver, and from childhood cancers, including recurrent and metastatic childhood liver tumours (hepatoblastoma and hepatocellular carcinoma). Importantly, these mutation catalogues were derived by orthogonal, non-duplex DNA sequencing and by independent variant calling. Tumour recurrences and metastases serve as reporters of past exposures, detectable by conventional bulk WGS (**Fig. 1A-B**), which has been the basis of the discovery of chemotherapy mutagenesis in the past^5^. To illustrate this point, we examined a recurrent hepatoblastoma in our cohort (PD62346) by conventional WGS, which demonstrated signatures of platinum agent exposure as well as the new signature in the tumour samples (**fig. S8A**). Prior to the resection of the recurrent hepatoblastoma, this child had received further platinum chemotherapy which therefore generated additional variants, as revealed by whole genome NanoSeq (**fig. S8B**).

In total we studied mutation data from five external cohorts: normal and cirrhotic liver conventional WGS data^39^; conventional WGS data from three different childhood cancer cohorts which included relapses of liver tumours^28,43^; and conventional whole exome sequencing data^44^ which included relapses and metastases of childhood liver tumours (**Fig. 4A**). We assessed published variant calls for mutational signatures, deriving consensus calls from *de novo* extraction and signature assignment. Note that amongst 1646 neoplasms, encompassing at least 60 different tumour types, there were 5 childhood liver tumour relapses that have been exposed to platinum agents during the treatment of the initial disease. *De novo* extraction and signature assignment singled out four of these (**Fig. 4B-C**) as harbouring the new signature. Mutational signatures of the fifth liver tumour relapse were unusual and indicative of sequencing artefacts (SBS53; **fig. S9**). In addition, *de novo* extraction and signature assignment demonstrated the new signature in pooled coding mutations from childhood liver metastasis and from recurrences (**Fig. 4C**). There was no evidence of the new signature in any other childhood cancer (including non-exposed liver tumours) or in adult cirrhotic liver samples (**Fig. 4C**). Overall, these independent mutation catalogues validate the new signature and corroborate that it seems to be specific to platinum-exposed liver tissues.

**Fig. 4.**
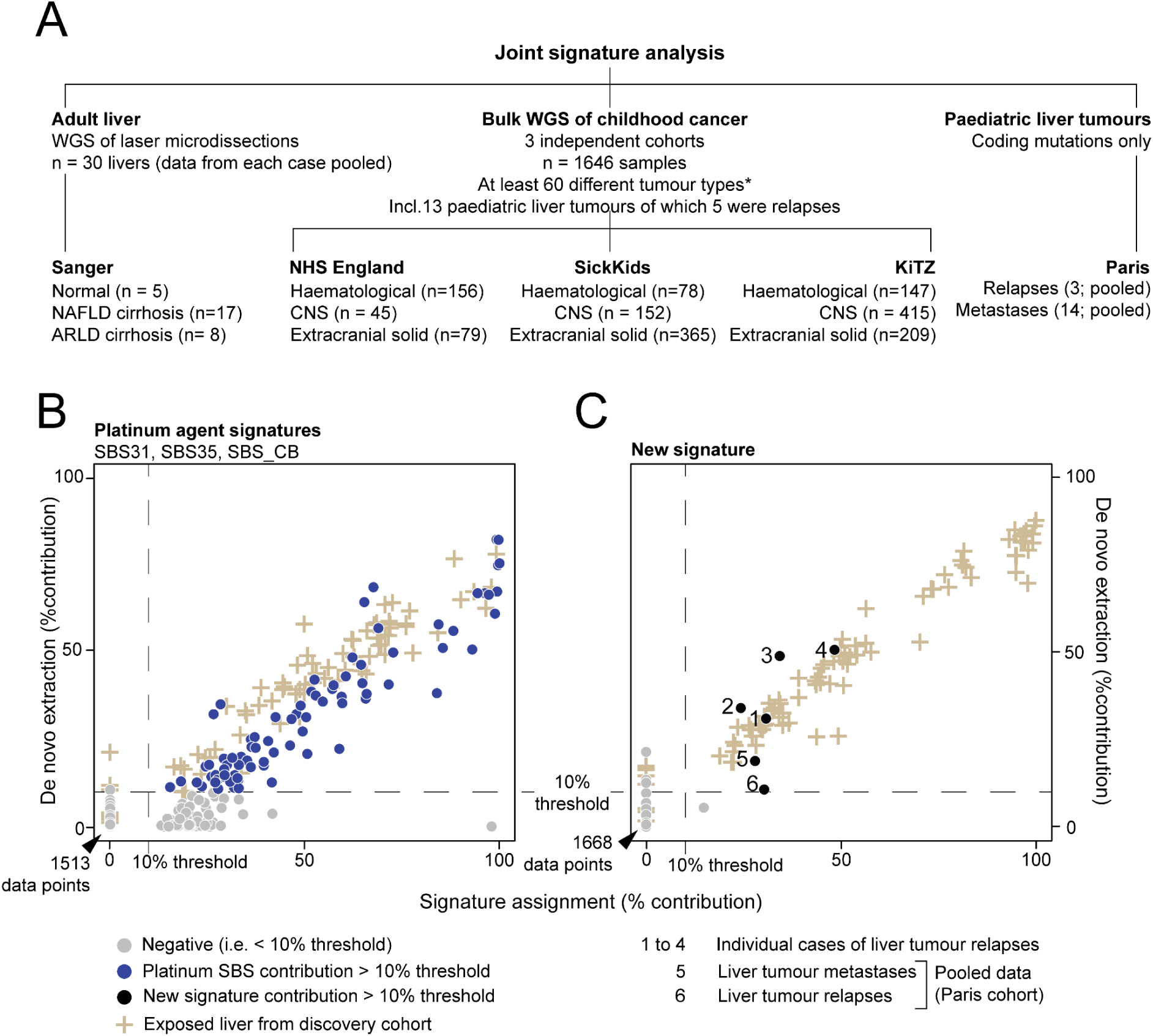
Signature extraction and assignment from mutation data of five relevant external cohorts. **(A)** Description of different data sets. Note that the childhood cancer data contained five recurrent childhood liver tumours from children who would have been treated with platinum agents prior to relapse. **(B)** Contribution of established platinum agent signatures to each sample. X-axis: contribution as per *de novo* extraction by mSigHdp. Y-axis: contribution as per signature assignment by SigProfilerAssignment (using signatures SBS1, SBS5, SBS40a, SBS31, SBS35, new signature, SBS_CB (carboplatin signature^5^). Dotted line in plot represents 10% threshold generally considered the level above which a signature result is considered to be robust. Symbols and colour coding as per legend. **(C)** Contribution of the new signature to each sample. Legends as per (C).

### Generation of pathogenic mutations

Targeted NanoSeq enabled us to examine whether chemotherapy mutagenesis generated variants that could have a functional impact (i.e. non-synonymous mutations) and that may be considered pathogenic in the context of mutation driven disease. We interrogated a total of 285 genes (**table S6**) including established cancer genes^38^ (n=231) and genes that have been found to harbour an excess of non-synonymous mutations in adult livers (n=7), some of which have been implicated as contributing to metabolic liver disease^39^. We divided coding variants into synonymous and non-synonymous. We then determined which non-synonymous variants in cancer genes would be considered pathogenic (“drivers”) in the context of cancer (for example, truncating variants in tumour suppressor genes or hotspot mutations in oncogenes). It is important to stress that a variant that would be annotated as a driver in a cancer may in fact be inconsequential in a normal cell (consider an inactivating mutation in one allele of a tumour suppressor gene when the state of the other allele in the same cell is unknown). Driver variants that we describe in normal tissues should, therefore, be considered as only potentially impactful. In exposed compared to non-exposed tissues, we found a significant enrichment of non-synonymous variants in general (p=2.2×10^-16^), of non-synonymous variants in genes related to metabolic liver disease (p=2.2×10^-16^), and of cancer driver mutations (p=2.2×10^-16^, Welch two-sided t-test for all three tests; **Fig. 5A**). Driver variants included tissue-relevant mutations, such as leukaemogenic events in blood or in normal liver, and exon 3 hotspot mutation of *CTNNB1* (the principal driver of hepatoblastoma). The enrichment of driver variants in exposed tissue could not be explained by technical variation in sequence coverage (**Fig. 5B**). It likely derives from the increased mutation burden associated with platinum exposure, with larger genes being more vulnerable to variation than smaller ones (**Fig. 5C**). Consistent with this proposition, we found that the new signature accounted for the majority of cancer driver variants (**Fig. 5D**). Furthermore, we did not find evidence that any particular cancer gene was under positive selection, as assessed from the ratio of non-synonymous over synonymous variants using an established statistical framework^45^. However, absence of selection should be viewed in the context of the short interval (weeks) between the last exposure and surgery. While platinum exposure generates a large number of non-synonymous variants, the time frame is likely too short for the selective advantage of mutant clones to emerge. One non-cancer gene that exhibited a significantly increased ratio of non-synonymous variants in all liver tissues, whether exposed or not, was the albumin gene, *ALB* (**fig. S10**). This increase has previously been observed in adult liver cancer and in non-transformed adult liver tissue^46^ and is of uncertain significance.

**Fig. 5.**
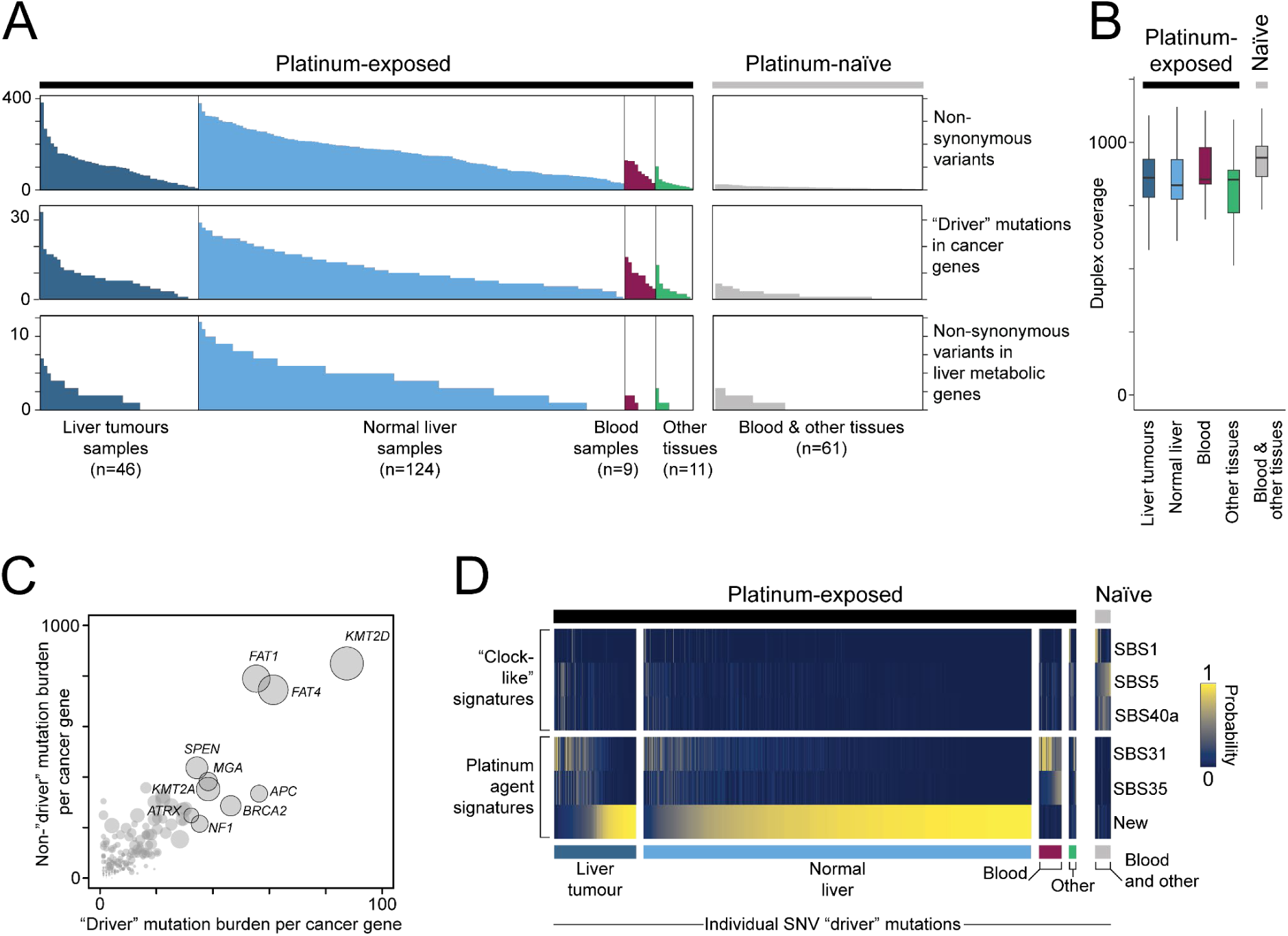
Annotonati of the functional potential of variants called by targeted NanoSeq. **(A)** Number of variants (Y-axis) per sample (X-axis; each bar represents one sample) in descending order and grouped as per labels. Top: all non-synonymous variants. Middle: variants that would be considered driver mutations in the context of cancer. Bottom: non-synonymous variants in genes in which mutations have been causally linked to adult metabolic liver disease^39^ (genes *ACVR2A*, *ALB*, *CIDEB*, *FOXO1*, *GPAM* and *TNRC6B*). There was a significant increase in variants comparing exposed with non-exposed tissues in all three panels (p=2.2*10-16; Welch two-sided t-test). Note that the order of the samples in the three panels are not the same. **(B)** Duplex coverage across groups. The duplex coverage is not higher in platinum-treated tissues than naïve tissues, and, therefore, does not cause the observed increase in non-synonymous variants and drivers in exposed tissues. **(C)** Correlation between non-driver mutation burden and potential driver mutation burden. Each dot represents a gene and the size of the dot indicates the coding sequencing length of the gene. In total, there were 2,022 potential driver variants and 21,623 non-driver mutations in 192 cancer genes (gene list constrained to those with at least one potential driver variant). Pearson correlation=0.85. Genes with a potential driver mutation burden above the 95th percentile are labelled by name. **(D)** Heatmap displaying the probability of potential cancer drivers (caused by a single base substitution; n=1,146) to have arisen due to a specific mutational signature. The variants are categorised by tissue type and their platinum exposure status as displayed in A. The majority of potential cancer drivers in platinum-exposed tissues were generated by the new signature or the known platinum signatures (SBS31 and SBS35).

## DISCUSSION

Almost all children who survive cancer develop late effects^17^. Some late effects have readily identifiable causes, such as sequelae from surgery. Others, such as secondary cancer or the early onset of degenerative diseases arise from chemotherapy exposure. The precise link between chemotherapy exposure and disease often remains elusive. Direct mutagenesis by cytotoxic agents represents a plausible molecular correlate of late effects. Our findings demonstrate that mutagenesis by carboplatin and cisplatin is pervasive in all paediatric normal tissues that we examined.

Our work resolves conflicting results of the past and, more importantly, reveals the full extent of mutagenesis: Platinum agents elevated the mutation burden of paediatric tissues to those seen in adults. Deploying targeted duplex sequencing, we were able to show a direct link of chemotherapy mutagenesis to the generation of potentially functional variants, such as cancer “driver” mutations in blood and liver. For example, in exposed normal liver amongst 105,414 molecules of DNA, we found 20,009 non-synonymous variants and 1,420 potentially pathogenic variants residing in cancer genes. This would suggest that within the millions of cells of a whole human liver, chemotherapy is likely to have generated a vast repertoire of potentially pathogenic mutations. Some of the diseases that survivors of childhood cancer experience in later life, such as second cancers and some metabolic liver diseases^17,18^, are caused by somatic mutations. It is conceivable therefore that an increase of potentially pathogenic variants in normal tissues of exposed children elevates the risk of developing mutation driven disease. If so, chemotherapy mutagenesis may represent a direct and quantifiable cause of late adverse effects from cancer treatment in childhood.

We made the remarkable discovery of liver specific mutagenesis. The significance of this finding does not lie in documenting a further mutational signature but in the discovery that body-wide (systemic) exposure to the same agents drove tissue-specific mutational processes. We were able to further corroborate this finding through re-examination of published cancer mutation catalogues, generated by independent sequencing and analysis pipelines. Tissue-specific mutagenesis has been observed with localised mutagens, such as UV light mutagenesis in skin^3,47^. It is, however, unprecedented that a systematic mutagen penetrating all tissues in the body causes a tissue-specific mutational signature. In the context of the function of the liver, it would seem intuitive to implicate hepatic metabolism in the generation of the new signature, akin to liver mutagenesis caused by metabolism of aflatoxins (carcinogens associated with adult liver cancer)^48,49^. Cisplatin and carboplatin undergo breakdown in the liver, where a series of reactions produce a circulating metabolite that is primarily secreted via the kidney into urine^50,51^. We did not find the new signature in exposed blood or other organs, indicating that if a metabolite were the culprit, it would probably be an intermediary that is contained within the liver. An implication of our discovery is that blood alone may not fully capture systemic mutagenic insults, which is particularly relevant to studies of chemotherapy mutagenesis that examine blood as a reporter of exposures. Furthermore, our discovery of differential mutagenesis raises the possibility that apparently distinctive mutational signatures may have been forged by the tissue context in which they were detected.

## Methods

### Ethics statement and sampling

Blood and tissue samples from children with liver tumours operated at the Institute of Liver Studies at King’s College Hospital were supplied via the King’s Paediatric Liver Research Biobank (REC ref: 23/WA/0059). Patients or guardians provided informed written consent for the storage and use of donated samples for research, with these samples subsequently released to this study as stipulated by the study protocol.

Foetal liver samples were obtained from the MRC–Wellcome Trust - funded Human Developmental Biology Resource (HDBR; http://www.hdbr.org^60^) with appropriate written consent and approval from the North East – Newcastle & North Tyneside 1 Research Ethics Committee. HDBR is regulated by the UK Human Tissue Authority (HTA; www.hta.gov.uk) and operates in accordance with the relevant HTA Codes of Practice. We enriched these foetal samples for hepatocytes through sorting of magnetic beads into CD45+ and CD45-fractions prior to nucleic acid extraction and sequencing.

All other human tissues were collected through studies approved by UK NHS research ethics committees. Patients or guardians provided informed written consent for participation in this study as stipulated by the study protocols. These studies have the following references: NHS National Research Ethics Service reference 16/EE/0394; 16/LO/0960.

### Nucleic acid extraction

Samples were stored fresh-frozen in −80°C until nucleic acid extraction according to standard methods using the Qiagen EZ1&EZ2 DNA Tissue kit for all tissues, except for blood where the Qiagen EZ1&2 DNA Blood kit was used.

### Conventional whole genome sequencing and filtering of variants

Short insert genomic libraries and flowcells were prepared followed by 150 bp paired-end sequencing clusters that were generated on Illumina HiSeq X. Resulting DNA sequences were aligned to the GRCh38 reference genome with the Burrows-Wheeler algorithm^52^.

Variants were called using extensively validated pipelines at our Institute: substitutions were called by CaVEMan^33^, indels by Pindel^53^, and copy-number changes by ASCAT^54^. CaVEMan and Pindel was run unmatched against an *in silico* human reference genome. Indel variants were filtered out if they failed Pindel’s filter or if they had a quality score below 300. Substitutions that passed CaVEMan’s filters were brought forward and mapping artefacts were removed by setting a threshold of ASMD ≥ 140, meaning the median alignment score of reads supporting a variant, and CLPM = 0, which requires that fewer than half of the reads were clipped. Germline substitutions and indels were filtered out by a one-sided exact binomial test as described previously^55^. Substitutions in diploid regions with consistently high or low number of reads across normal samples from the same individual were removed. Next, all remaining substitutions and indels in samples from the same individual were recounted twice: once where we set a threshold of read mapping quality=30 and base quality=25 (high-quality reads), and once without applying any thresholds for mapping and base quality (total reads). Variants were included if the fraction of high-quality reads over total reads was greater than 75%. To separate true substitutions from noise, we employed a beta-binomial model as described previously^55^ where the locus-specific error rate is calculated using blood samples from healthy individuals as the reference. The resulting P values were corrected for multiple testing with the Benjamini–Hochberg procedure and a cut-off was set at q < 0.01. Additionally, artefactual substitutions were removed if they were within 10 base pairs of where an indel had been called. Indels were removed if any indel had been called at the same position in the matched blood sample, and artefacts were also removed if any failed indel (indicative of a position with worse quality or mapping issues) had been called at that position in any other sample. Finally, all variants were manually inspected with jBrowse^56^ to remove artefacts.

### NanoSeq – library preparation, sequencing and variant calling

Whole genome NanoSeq libraries were prepared as previously described^31^ and sequenced to ∼30 duplex coverage on Illumina NovaSeq 6000. Targeted NanoSeq libraries were prepared according to Lawson et al.^32^, with 2 rounds of pulldown using a custom bait set (TWIST) consisting of 285 genes (∼1 MB in size, **table S6**) and sequenced on Illumina NovaSeq 6000 (median duplex coverage ∼860 per sample). Sequenced DNA reads were aligned to GRCh37 (hs37d5 build) for targeted NanoSeq data and GRCh38 for whole genome NanoSeq using the Burrows-Wheeler algorithm. Variants were called with the established NanoSeq pipeline as previously described^31^ (https://github.com/cancerit/NanoSeq). Conventional WGS of blood samples (median coverage of 88x per sample) from the same individual was used as the matched normal in the variant-calling pipeline for whole genome NanoSeq to remove germline variants. Matching targeted NanoSeq of blood samples was used to filter out germline variants in targeted NanoSeq data. If a matching blood sample was unavailable, we utilised the high sequencing depth and captured polyclonality of targeted NanoSeq to filter out variants with a variant allele frequency >10% (including germline variants) instead of using a matched sample.

### NanoSeq – normalisation of mutation burden and trinucleotide substitution profiles

Mutation rates are heavily affected by the trinucleotide context and we therefore normalised the obtained NanoSeq substitution burdens and trinucleotide frequencies to account for differences between the regions sequenced with NanoSeq and the genome-wide background, as described previously^31^. The normalised trinucleotide frequencies and normalised substitution burdens are used throughout the paper. The mutation burden is multiplied by the size of the human diploid genome (∼6 billion base pairs) to obtain the number of mutations per diploid genome, which is compared to the number of mutations obtained by conventional WGS.

### Filtering of targeted NanoSeq data

No samples for which data were generated were excluded from targeted NanoSeq analyses. Artefacts in targeted NanoSeq data may occur towards the start and end of reads, often because of germline indels at the edge of a read which can be miscalled as a substitution. To remove these, a Kolmogorov-Smirnov (KS) test on read position was applied across a large targeted NanoSeq dataset. Mutations were discarded if the KS test q value was less than 0.05 and more than 50% of the mutations were within the first or last 10 bp of callable reads. A subset of called mutations that had not been discarded were reviewed manually on jBrowse^56^ and were consistent with true mutations.

### Driver annotation from targeted NanoSeq data based on prior knowledge

Variants were annotated with the dndscv package^45^ and were annotated as driver mutations if they fell into one of two categories: truncating mutations in tumour suppressor genes, or nonsynonymous mutations in known hotspots (positions within genes that are excessively mutated).

For the annotation of truncating mutations in tumour suppressor genes, four lists of tumour suppressor genes were defined: the COSMIC Cancer Gene Census^38^, and lists based on curation of the literature for each of liver cancers, blood cancers, and Wilms tumours. The Cancer Gene Census should capture common pancancer genes, while tissue-specific lists were chosen based on the makeup of our dataset. A truncating (nonsense, frameshift, or splice site) mutation in any tumour suppressor in any of these lists was taken forwards as a candidate driver mutation.

For the annotation of hotspot mutations, four lists of hotspot positions were generated using COSMIC data: across all cancer, and specifically in each of liver cancers, blood cancers, and Wilms tumours. Within each group, for every gene, the number of substitutions in every codon of the gene was counted. For each gene, the median number of mutations across all codons was calculated. If a codon had greater than four times the median plus one (one is added for cases where the median number of mutations is zero), or if it had more than 15 mutations in total, that codon was counted as a hotspot codon. Any nonsynonymous substitution in a hotspot codon was counted as a candidate driver mutation.

### Selection analyses from targeted NanoSeq data

Selection analyses were carried out using the package dndscv^45^ for the following sample groups: all samples together; normal liver with and without platinum exposure; all normal tissues with and without platinum exposure; blood with and without platinum exposure; and hepatoblastoma tumours with and without platinum exposure. For every group, dndscv was run with the following parameters, where the gene list included 275 protein coding genes included in the baitset:

dnds_output=dndscv(dndsin_input_mutations, max_muts_per_gene_per_sample = Inf, max_coding_muts_per_sample = Inf, gene_list=genelist, onesided = T, dc=duplex_coverage_vector, outmats=T).

Genes were considered to be under positive selection if their positive selection q value across all mutation classes was <0.05.

### Targeted NanoSeq – probability of a variant arising from a specific mutational signature

The probability of a certain substitution arising from a specific signature was calculated as previously described^26^ where the weighted fraction of the signature contribution to the sample is multiplied by the mutational probability of the signature to cause the substitution in that specific trinucleotide context:

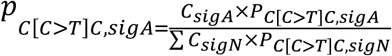

I.e, the odds of signature A causing a C[C>T]C mutation is calculated by taking the contribution of signature A (C,sigA) to the sample (as defined by signature assignment or *de novo* extraction followed by decomposition) multiplied by the probability (P) that signature A generates a C[C>T]C change (percentage of SBS for a C[C>T]C change in signature A in COSMIC SBS96 reference database). This is then normalised by the sum of these products for all signatures (N: number of signatures) detected in the sample of interest. As a result, the probability for each signature to have caused the substitution of interest will range between 0 and 1 with the highest score representing the most likely signature.

### *De novo* mutational signature extraction and decomposition

SBS96, ID83 and DBS72 *de novo* signature extraction was performed by two methods. For both methods, *de novo* signatures with a contribution lower than 10% to a sample were considered unassigned. Method 1) SigProfilerExtractor^37^ was run using default settings and a maximum of 20 signatures to be extracted. The suggested solution of the number of extracted signatures by the algorithm was used and decomposition into COSMIC reference version 3.4 was performed by the SigProfilerExtractor tool. Method 2) mSigHdp^36^ was run using the recommended values for gamma.beta (20 for SBS, 50 for indels and DBS extraction), number of burn-in (100,000 for SBS and DBS, 30,000 for indels) and number of chains/iterations (20 independent chains, 200 Gibbs samples per chain and with 100 iterations between collected samples). Three iterations of concentration parameters were performed after each iteration. For the mSigHdp extraction of external datasets (Fig. 4), we set downsampling to 3000 (as recommended) to reduce computational demand for processing this highly mutated dataset and also reduced the number of burn-in to 10,000. Signatures extracted by mSigHdp were decomposed into linear mixtures of COSMIC v3.4 reference signatures using an expectation maximization algorithm. Cosine similarity between the extracted signature and the reconstructed signature based on the decompositions were calculated both for the SigProfiler and mSigHdp extractions. Extracted signatures with a cosine similarity >=0.9 was considered to be combinations of previously reported signatures, whereas a cosine similarity <0.9 was considered insufficient to explain the extracted signature and thus suggestive of a novel extracted signature.

SigProfilerExtractor^37^ was used to extract signatures based on the SBS288 classification (an extension of the SBS96 classification where it also considers the strand orientation of the mutation) to check for transcriptional strand bias of the new signature.

*De novo* signature extraction of the targeted NanoSeq data for the validation cohort was performed to assess the new signature as extracted by targeted NanoSeq data, which was reassuringly very similar (cosine similarity 0.97) to the new signature as extracted from WGS NanoSeq discovery cohort. The extraction was also performed to get SBS signature proportions that were used for subsequent calculations of driver probability (as described above). The validation cohort contains an overwhelmingly majority of platinum-exposed samples and mutations and we therefore supplemented this cohort with 120 blood samples (**table S8**) from platinum-naïve children with Wilms tumour to extract the clock-like signatures.

### Mutational signature assignment

We used two different methods for signature assignment of SBS1, SBS5, SBS40a, SBS31, SBS35 (all from COSMIC SBS96 reference version 3.4), our new signature as extracted by mSigHdp (from WGS NanoSeq or targeted NanoSeq depending on the sequencing type of the samples), and SBS-Carboplatin (as extracted previously^5^ using SignatureAnalyzer). The first method, SigProfilerAssignment^40^, was run with the function cosmic_fit using default settings restricted to the aforementioned signatures. mSigAct^41^ was applied using the function SparseAssignActivity, with its recommended settings and the p-value threshold was set to 0.05 divided by the number of signatures included for assignment. Signatures with a contribution lower than 10% to a sample were considered unassigned.

## Data availability

De-identified patient-level information and variant calls are provided in Supplementary Tables. Sequencing data are available through the European Genome-Phenome Archive (EGA), accession code pending. External datasets analysed in this study are either published ^28,39,43,44^ or available upon request from Adam Shlien (SickKids WGS study).

## ACKNOWLEDGEMENTS

We thank Sir Michael Stratton (Wellcome Sanger Institute) and Aidan Maartens (science writer, Wellcome Sanger Institute) for critical review of this manuscript, and liver transplant coordinators and paediatric clinical nurse specialists at King’s College Hospital NHS Foundation Trust. We also thank the funders of this study: The Wellcome Trust (personal fellowship to S. Behjati, reference 223135/Z/21/Z and to A. Hodder; institutional grant to the Wellcome Sanger Institute, reference 220540/Z/20/A), The Wenner-Gren Foundations (personal fellowship to A. Wenger), The Children’s Kidney Care Fund (University of Cambridge; support of J. Kennedy), FAPESP 2023/12984-7 (personal fellowship to F.L.T. Silva), King’s Health Partners Centre for Translational Medicine (personal fellowship to J.J. Sun), The Little Princess Trust, NIHR GOSH Biomedical Research Centre and NIHR Cambridge Biomedical Research Centre (NIHR203312). The funders had no role in study design, data collection and analysis, decision to publish or preparation of the paper. The views expressed are those of the authors and not necessarily those of the NHS, the NIHR or the Department of Health. Foetal liver samples were obtained from the MRC–Wellcome Trust - funded Human Developmental Biology Resource. We are indebted to the children and families who participated in this study.

## AUTHOR CONTRIBUTIONS

Conceptualization: AW, FJR, SB

Data curation: MD, ML, TO, CP, TD, MH, JJS, SA, WJ, CT, AD, VJ, NH, FJR, SB

Formal analysis: AW, HLS, SB

Funding acquisition: SB

Investigation: AW, HLS, MCC, MD, YZ, FJR, SB

Methodology: AW, SB

Project administration: SB

Resources: MD, ML, ARJL, FA, PAN, TDT, TO, TRWO, JK, AH, NDA, FLTS, MKT, TD, MH, KS, IM, LP, JCT, CT, NH, AS, FJR, SB

Software: AW, HLS, ARJL, FA

Supervision: ARJL, FA, IM, THHC, FJR, SB

Validation: AW, ML, AS, SB

Visualization: AW, HLS, SB

Writing – original draft: AW, SB

Writing – review & editing: AW, ARJL, TDT, THHC, FJR, SB

