## Supplementary material for "Extensive and differential platinum chemotherapy mutagenesis in children": Supp figures

**The PDF file includes:**

Figs. S1 to S10

**Other Supplementary Materials for this manuscript include the following:**

Tables S1 to S8

A

| | PD42751 | PD64735 | PD57225 | PD57226 | PD59132 | PD62345 |
|---|---|---|---|---|---|---|
| 11 p | LOI | LOH | LOH | | LOH | LOH |
| *CTNNB1* | | | G34V | ess. splice | | |
| *GNA11* | | | | R183H | | |

Driver (shaded) / No (unshaded)

B

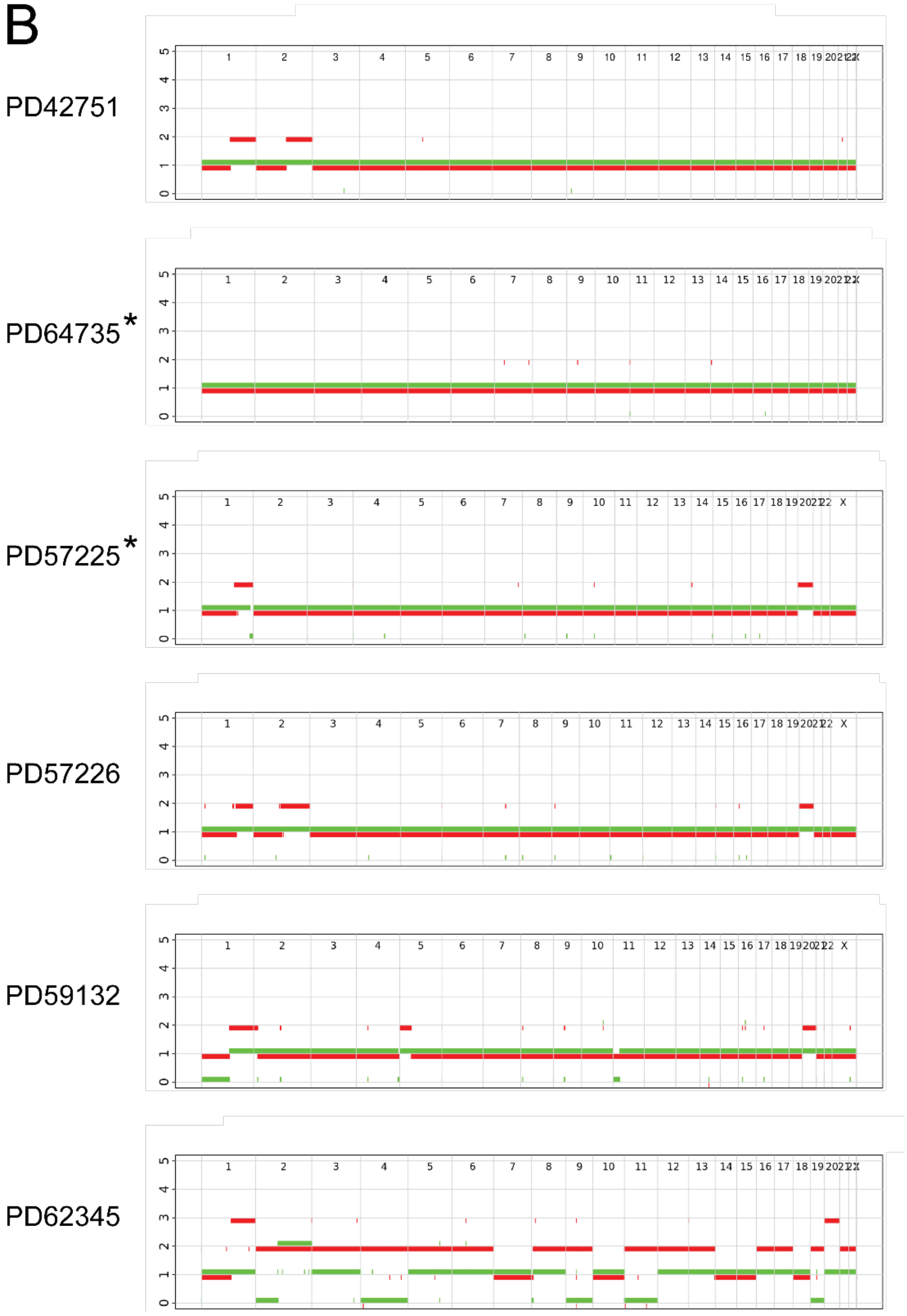

**Fig. S1. Driver variants and copy-number alterations in hepatoblastoma tumours.**
**(A)** Drivers (based on conventional WGS) in the tumours in the discovery cohort include 11p loss of imprinting (LOI) and loss of heterozygosity (LOH), as well as somatic substitutions and small insertions/deletions (indels).
**(B)** ASCAT copy-number plots (based on conventional WGS) of one representative tumour biopsy per case in the discovery cohort. A normal diploid genome without any copy-number aberrations has one copy each for every somatic chromosome (green and red line at 1). The tumours have common alterations in hepatoblastoma such as 11p LOH, gain of 1q, gain of 2q and gain of chromosome 20. * Mosaic 11p LOH is present in the germline and prevents 11p LOH from being detected in the tumour sample since the copy-number calling is run against the blood sample.

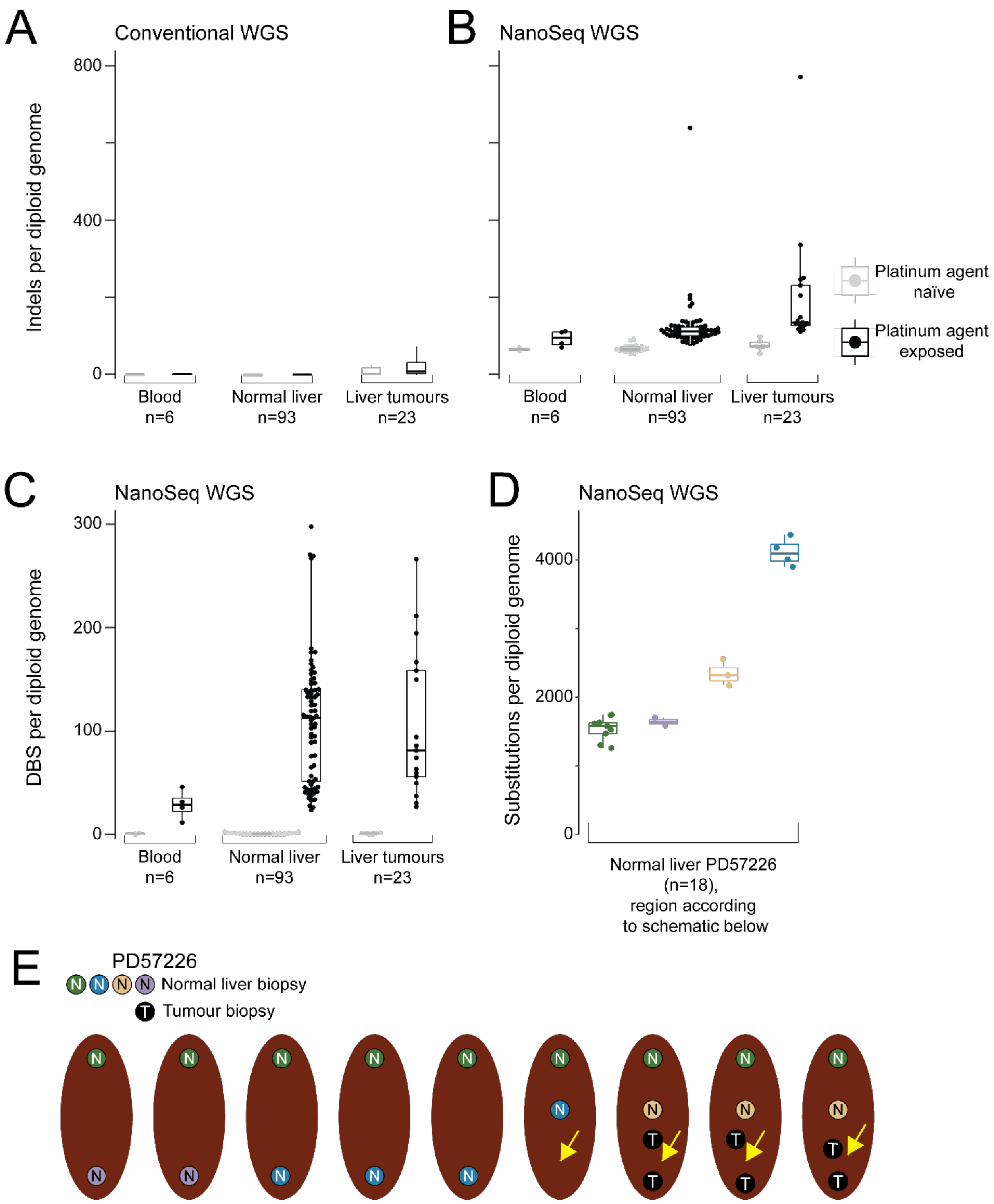


**Fig. S2.Indel and double base substitution burden in the discovery cohort.**
**(A)** Indel burden by conventional whole genome sequencing (WGS). X-axis: different tissues divided based on platinum exposure as shown in legend in (B). Y-axis: number of indels per diploid genome.
**(B)** Mutation burden by WGS NanoSeq. Axes and horizontal lines as per (A).

**(C)** Double base substitution (DBS) burden by whole genome NanoSeq. X-axis: different tissues divided based on platinum exposure as shown in legend in (B). Y-axis: number of DBS per diploid genome.
**(D)** Single base substitution burden by whole genome NanoSeq for PD57226 shows regional variation in normal liver samples coloured according to their sampling location in (E).
**(E)** Sampling schematic of PD57226. Normal liver (N) and tumour (T) tissue. Yellow arrows delineate the tumour.

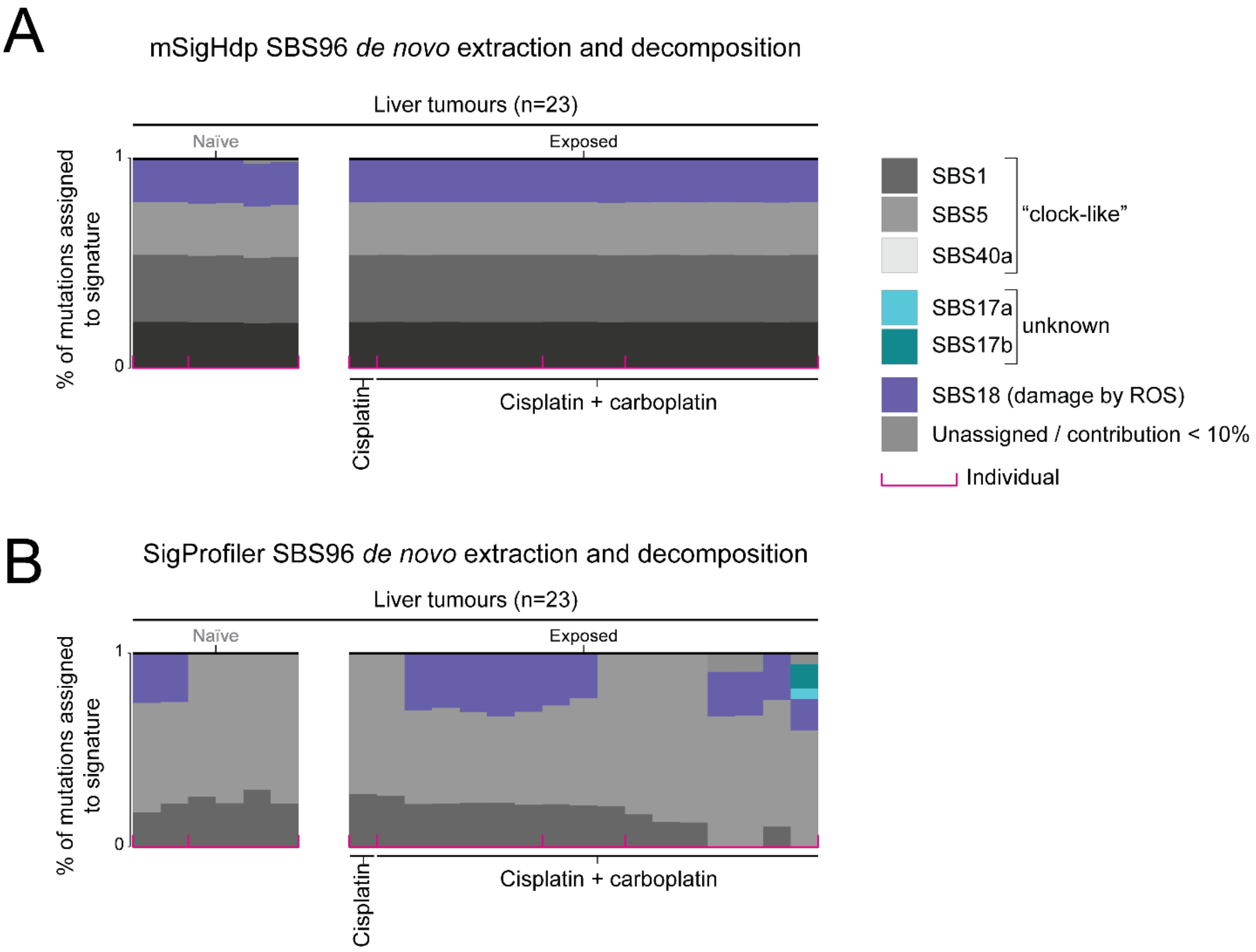


**Fig. S3. *De novo* signature extraction of single base substitutions called by conventional WGS.**

**(A)** mSigHdp *de novo* extraction and subsequent decomposition into known COSMIC SBS96 signatures of single base substitutions from liver tumours based on conventional WGS. Contribution of signatures to each sample. X-axis: Each stacked bar plot represents a tissue sample as per labelling. Y-axis: % of mutations assigned to signature. Pink square brackets delineate individuals. Colours represent mutational signatures as per legend.

**(B)** SigProfiler *de novo* extraction and subsequent decomposition into known COSMIC SBS96 signatures of substitutions from liver tumours based on conventional WGS. Same setup of plot as in (A). Note that the new mutational signature was not detected by conventional WGS and neither was any of the known platinum signatures (SBS31 and SBS35).

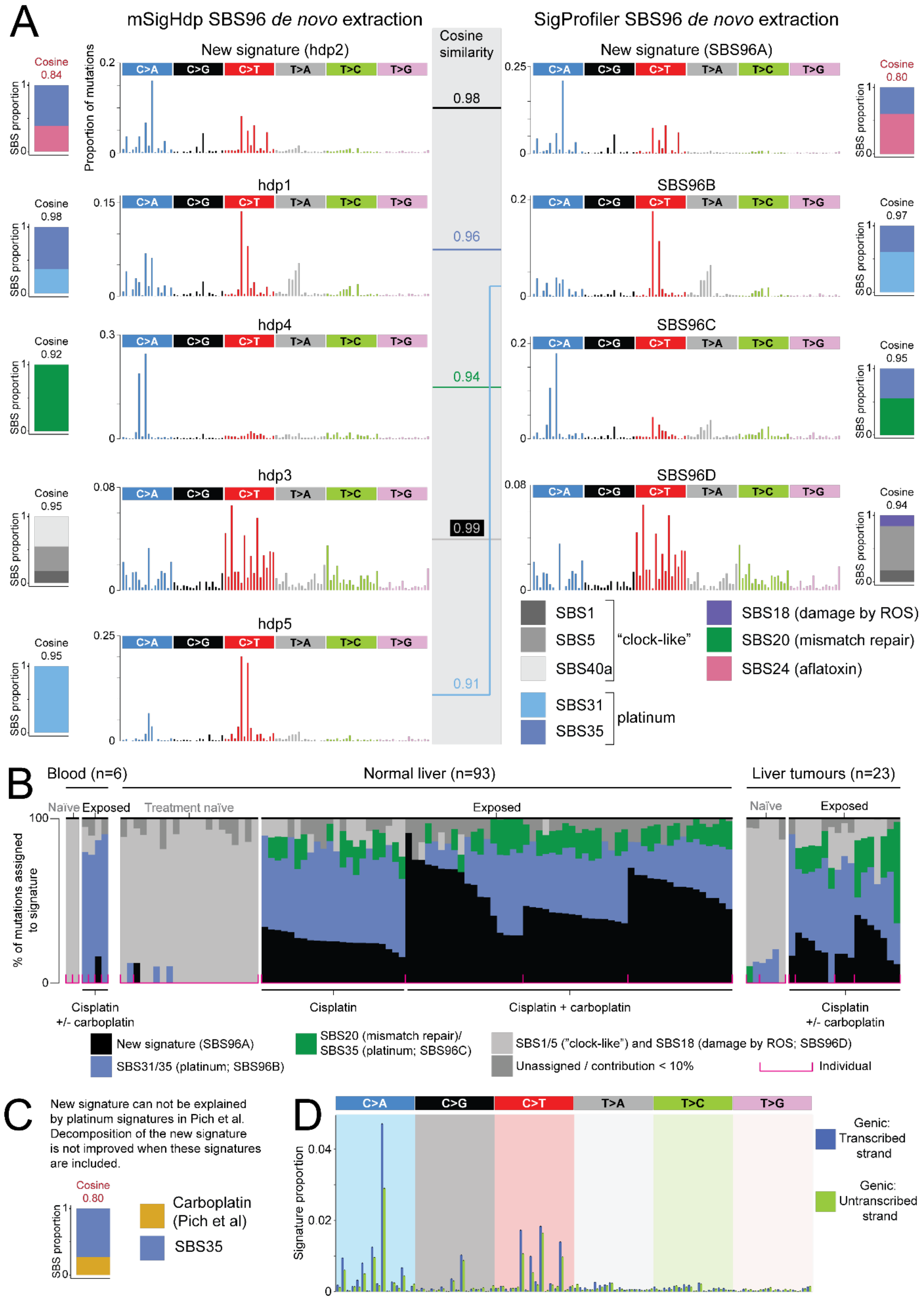
A
mSigHdp SBS96 de novo extraction
SigProfiler SBS96 de novo extraction
Cosine similarity
New signature (hdp2)
New signature (SBS96A)
hdp1
SBS96B
hdp4
SBS96C
hdp3
SBS96D
hdp5
0.98
0.96
0.94
0.99
0.91
Proportion of mutations
SBS proportion
C>A C>G C>T T>A T>C T>G
SBS1
SBS5
SBS40a
"clock-like"
SBS31
SBS35
platinum
SBS18 (damage by ROS)
SBS20 (mismatch repair)
SBS24 (aflatoxin)
B
Blood (n=6)
Normal liver (n=93)
Liver tumours (n=23)
Naïve
Exposed
Treatment naïve
% of mutations assigned to signature
Cisplatin +/- carboplatin
Cisplatin
Cisplatin + carboplatin
New signature (SBS96A)
SBS31/35 (platinum; SBS96B)
SBS20 (mismatch repair)/ SBS35 (platinum; SBS96C)
SBS1/5 ("clock-like") and SBS18 (damage by ROS; SBS96D)
Unassigned / contribution < 10%
Individual
C
New signature can not be explained by platinum signatures in Pich et al. Decomposition of the new signature is not improved when these signatures are included.
Cosine 0.80
Carboplatin (Pich et al)
SBS35
D
Signature proportion
Genic: Transcribed strand
Genic: Untranscribed strand

**Fig. S4. *De novo* signature extraction of single base substitutions called by whole genome NanoSeq.**

**(A)** The five extracted *de novo* SBS96 signatures by mSigHdp (second column) based on whole genome NanoSeq data of the discovery cohort, and the known SBS signatures they decompose into (first column; the reconstructed cosine similarity is stated on top of the stacked bar plot). Decompositions with a reconstructed cosine similarity below 0.9 (marked in red) were considered as insufficient to explain the extracted signature. Colours of SBS signatures according to legend. The third column (grey box) shows the cosine similarity between the *de novo* signatures extracted by mSigHdp and SigProfiler (fourth column). The decomposition of the SigProfiler extracted signatures are shown to the far right (fifth column). The signatures extracted by the two methods were similar.

**(B)** Contribution of signatures to each sample. X-axis: Each stacked bar plot represents a tissue sample as per labelling. Y-axis: % of mutations assigned to signature. Pink square brackets delineate individuals. Colours represent mutational signatures as per legend. The SigProfiler extraction is shown here. Similar results were obtained by mSigHdp (Fig. 2E). Note that the new signature was only evident in platinum-exposed livers whereas conventional platinum signatures were pervasive across all platinum-exposed tissues.

**(C)** We included platinum signatures as described in the literature (*5*), but the new signature could not be decomposed into those and they do not explain the new signature.

**(D)** The new signature displays transcriptional strand bias.

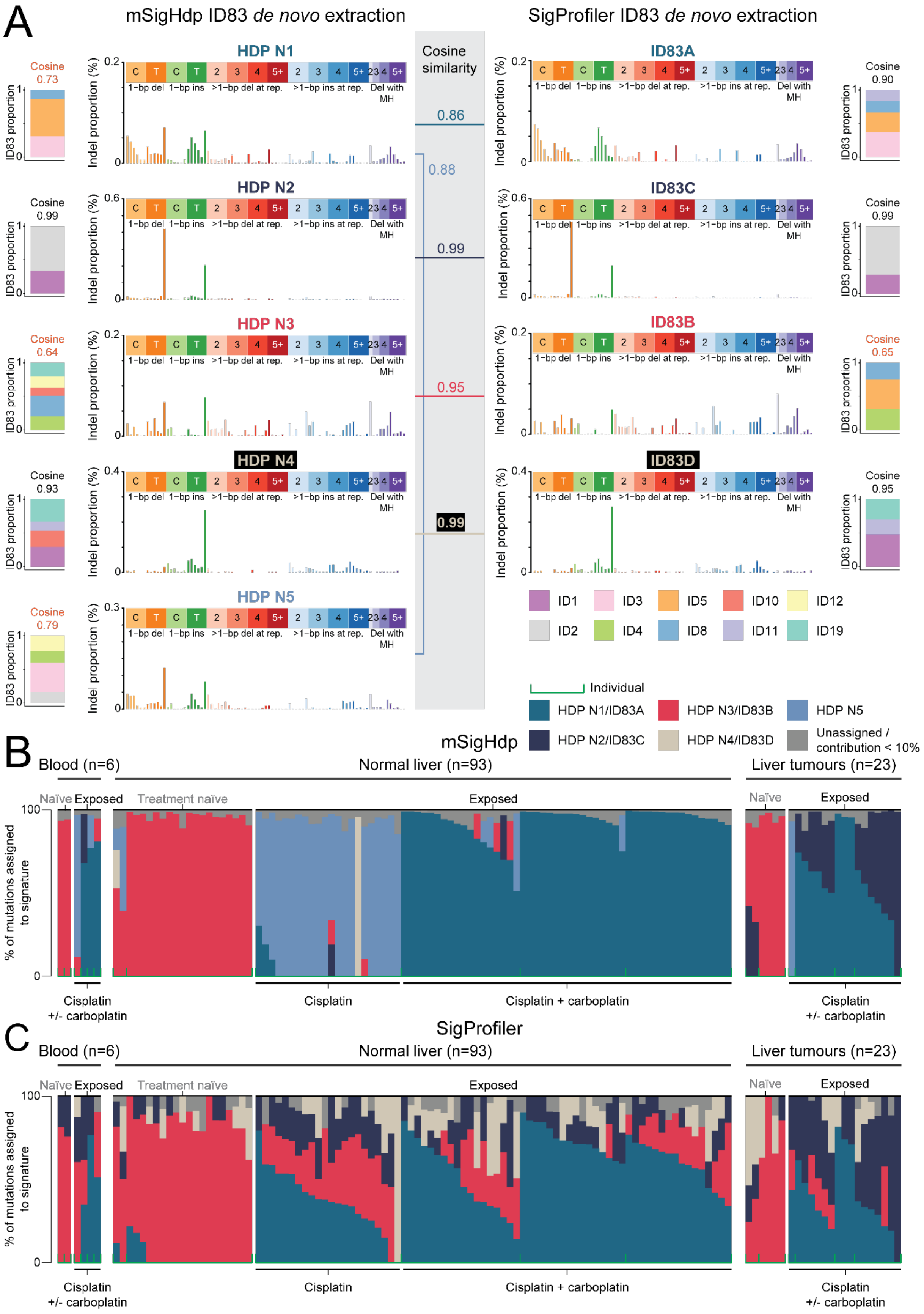

A
mSigHdp ID83 de novo extraction
SigProfiler ID83 de novo extraction
Cosine similarity
HDP N1
HDP N2
HDP N3
HDP N4
HDP N5
ID83A
ID83C
ID83B
ID83D
0.86
0.88
0.99
0.95
0.99
Cosine 0.73
Cosine 0.99
Cosine 0.64
Cosine 0.93
Cosine 0.79
Cosine 0.90
Cosine 0.99
Cosine 0.65
Cosine 0.95
ID83 proportion
Indel proportion (%)
1-bp del
1-bp ins
>1-bp del at rep.
>1-bp ins at rep.
Del with MH
ID1
ID3
ID5
ID10
ID12
ID2
ID4
ID8
ID11
ID19
Individual
HDP N1/ID83A
HDP N3/ID83B
HDP N5
HDP N2/ID83C
HDP N4/ID83D
Unassigned / contribution < 10%
B
mSigHdp
Blood (n=6)
Normal liver (n=93)
Liver tumours (n=23)
Naïve
Exposed
Treatment naïve
% of mutations assigned to signature
Cisplatin +/- carboplatin
Cisplatin
Cisplatin + carboplatin
C
SigProfiler

**Fig. S5. *De novo* signature extraction of insertion/deletions called by whole genome NanoSeq.**

**(A)** The five extracted *de novo* indel signatures by mSigHdp (second column) based on whole genome NanoSeq data of the discovery cohort and the known ID83 signatures they decompose into (first column; the reconstructed cosine similarity is stated on top of the stacked bar plot). Decompositions with a reconstructed cosine similarity below 0.9 (marked in orange) were considered as insufficient to explain the extracted signature. Colours of ID signatures according to legend. The third column (grey box) shows the cosine similarity between the *de novo* signatures extracted by mSigHdp and SigProfiler (fourth column). The decomposition of the SigProfiler extracted signatures are shown to the far right (fifth column).

**(B)** Contribution of signatures to each sample. X-axis: Each stacked bar plot represents a tissue sample as per labelling. Y-axis: % of mutations assigned to signature. Green square brackets delineate individuals. Colours represent mutational signatures as per legend. The mSigHdp extraction is shown here.

**(C)** Similar results were obtained by the SigProfiler extraction.

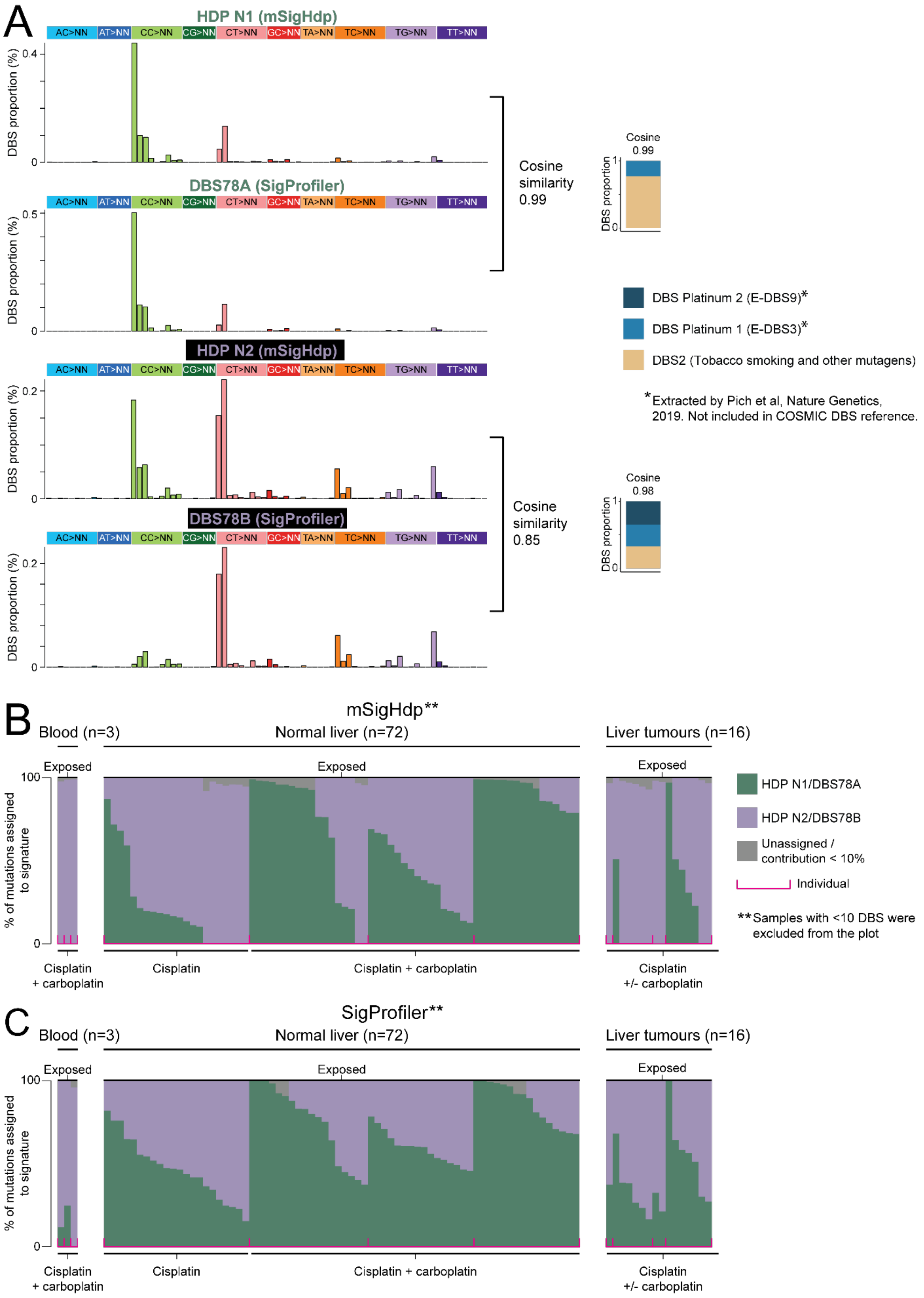
A
HDP N1 (mSigHdp)
AC>NN AT>NN CC>NN CG>NN CT>NN GC>NN TA>NN TC>NN TG>NN TT>NN
DBS proportion (%)
DBS78A (SigProfiler)
Cosine similarity 0.99
Cosine 0.99
DBS proportion
HDP N2 (mSigHdp)
DBS78B (SigProfiler)
Cosine similarity 0.85
Cosine 0.98
DBS Platinum 2 (E-DBS9)*
DBS Platinum 1 (E-DBS3)*
DBS2 (Tobacco smoking and other mutagens)
*Extracted by Pich et al, Nature Genetics, 2019. Not included in COSMIC DBS reference.
B
mSigHdp**
Blood (n=3)
Normal liver (n=72)
Liver tumours (n=16)
Exposed
% of mutations assigned to signature
HDP N1/DBS78A
HDP N2/DBS78B
Unassigned / contribution < 10%
Individual
**Samples with <10 DBS were excluded from the plot
Cisplatin + carboplatin
Cisplatin
Cisplatin + carboplatin
Cisplatin +/- carboplatin
C
SigProfiler**
Blood (n=3)
Normal liver (n=72)
Liver tumours (n=16)
Exposed
% of mutations assigned to signature
Cisplatin + carboplatin
Cisplatin
Cisplatin + carboplatin
Cisplatin +/- carboplatin

**Fig. S6. *De novo* signature extraction of double base substitutions called by whole genome NanoSeq.**

**(A)** The two extracted *de novo* double base substitution (DBS) signatures by mSigHdp and SigProfiler (left) based on whole genome NanoSeq data of the discovery cohort. One of the extracted signatures is extremely similar across the two methods (cosine similarity 0.99) whereas the other signature differs in terms of CC>NN mutations, but is otherwise similar between the two methods and thus visualised in the same colour in (B) and (C). The extracted signatures are best explained by using DBS2 (included in the COSMIC reference database) and two platinum signatures extracted by Pich et al. (*5*) (right). The platinum signatures extracted by Pich et al. are a refinement of the platinum signature DBS5 present in the COSMIC reference database. The reconstructed cosine similarity of the decomposition of the mSigHdp extracted signatures is stated on top of the stacked bar plot. Colours of DBS signatures according to legend.

**(B)** Contribution of signatures to each sample. X-axis: Each stacked bar plot represents a tissue sample as per labelling. Y-axis: % of mutations assigned to signature. Pink square brackets delineate individuals. Colours represent mutational signatures as per legend. Samples with fewer than 10 DBS were excluded from (B) and (C). The mSigHdp extraction is shown here. Note that signature HDP N1 was only present in platinum-exposed normal liver and tumours, but absent from platinum-exposed blood.

**(C)** Similar results were obtained by the SigProfiler extraction.

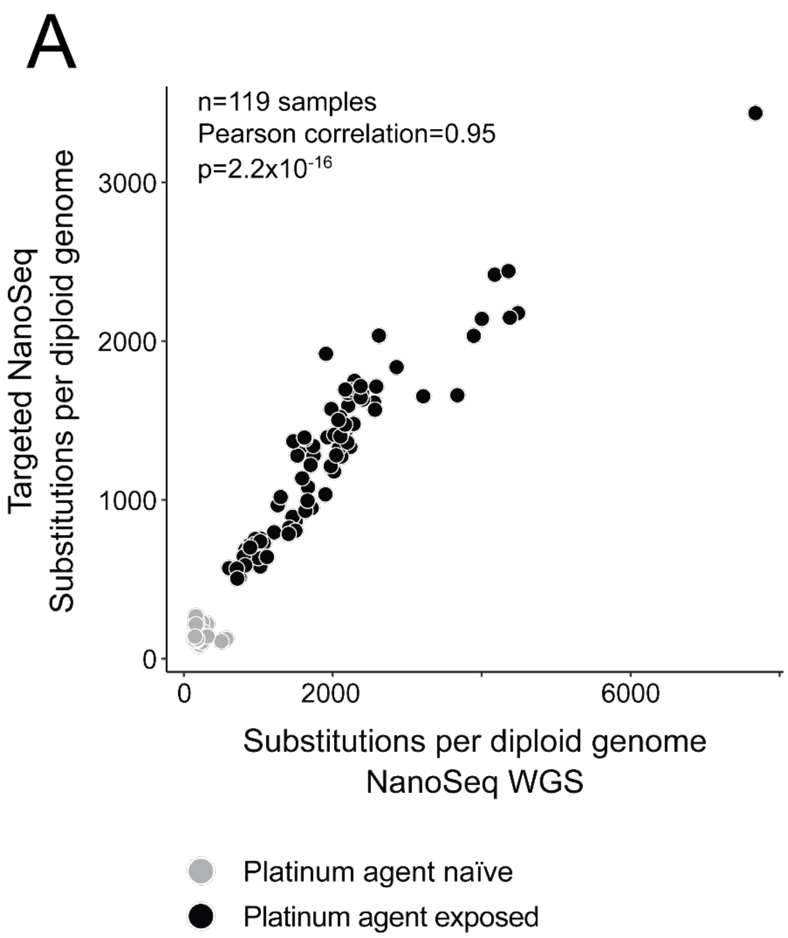


**Fig. S7. Single base substitution burden in targeted versus whole genome NanoSeq.**
**(A)** Mutation burden by targeted NanoSeq (y-axis) versus mutation burden by whole genome NanoSeq (x-axis) shows a high correlation (Pearson correlation=0.95, $p=2.2x10^{-16}$). Platinum-exposed samples are coloured in black and platinum-naïve in grey.

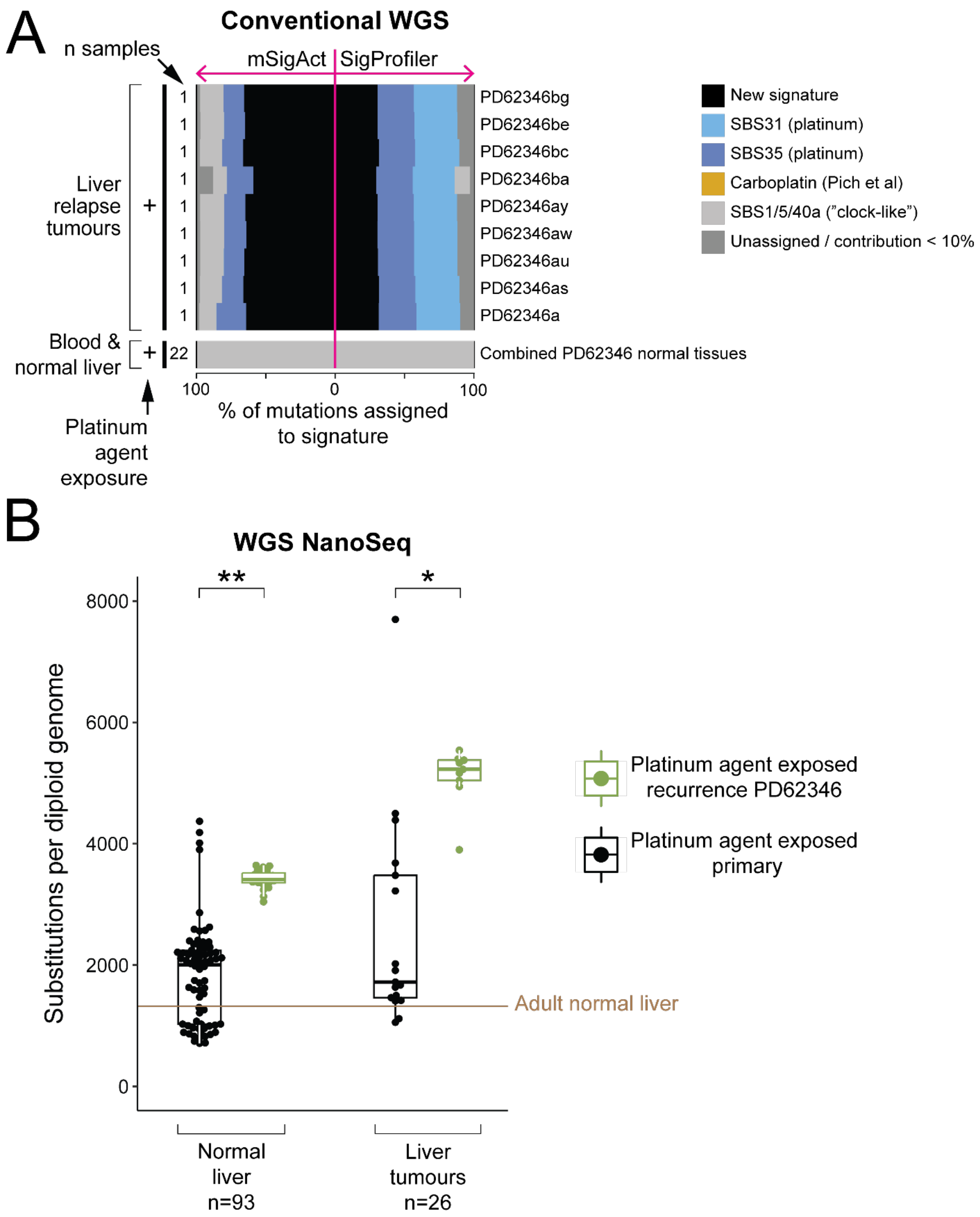


**Fig. S8. Signature assignment and mutation burden in a recurrent hepatoblastoma.** **(A)** Contribution of signatures to each sample based on conventional whole genome sequencing (WGS) of the recurrent liver tumour PD62346. X-axis: % of mutations assigned to signatures with each signature represented by a different colour. Stacked bar plots to the left of the pink line are results from mSigAct whereas stacked bar plots to the right are results from SigProfilerAssignment. Y-axis: Each stacked bar plot represents a tissue sample as per labelling. **(B)** Mutation burden by whole genome NanoSeq for platinum-treated normal liver and tumours respectively divided into all primary platinum-exposed primary hepatoblastoma (black) vs all platinum-exposed recurrent hepatoblastoma samples (PD62346; green). Samples from the recurrence have significantly higher mutation burden than samples from primary cases. The brown horizontal line represents mutation burden observed in normal adult liver (*34*). * $p=1.7x10^{-5}$, ** $p=2.2x10^{-16}$.

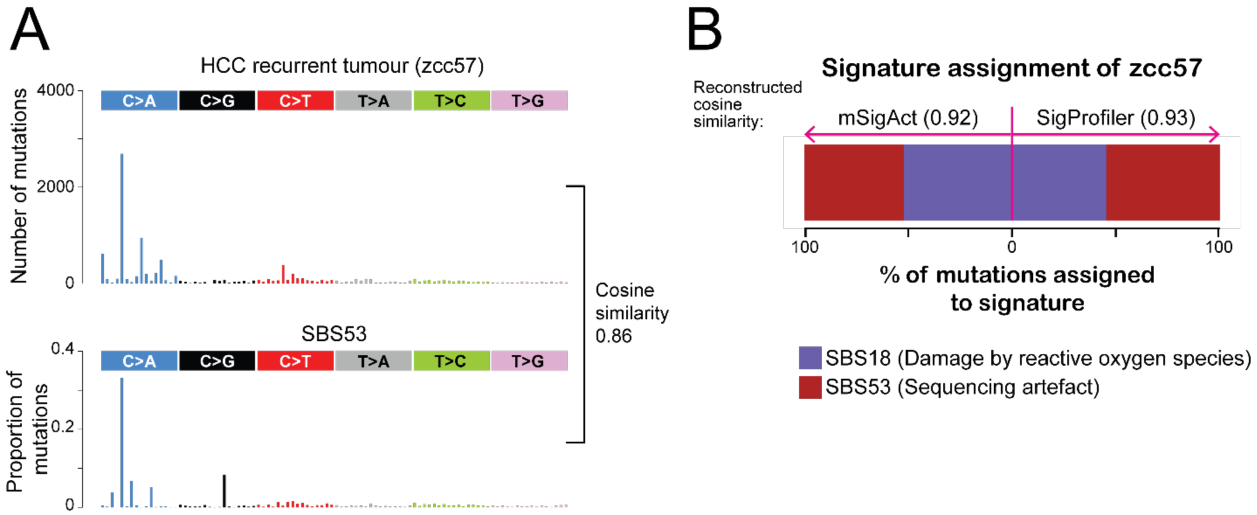


**Fig. S9. Mutational spectra and signature assignment of a recurrent hepatocellular carcinoma.**

**(A)** Trinucleotide spectra for zcc57 (top), a platinum-treated recurrent hepatocellular carcinoma tumour, which showed no signal of the new mutational signature. It displays a spectrum similar to SBS53 (bottom; cosine similarity 0.86), which is a sequencing artefact.

**(B)** Signature assignment by mSigAct (left of the pink vertical line) and SigProfilerAssignment (right of the pink vertical line) both show that sequencing artefacts (SBS53) are present in the zcc57 tumour sample, together with SBS18. The cosine similarity from the assignments compared to the original spectra of the tumour was high (0.92 and 0.93 respectively), indicating a reliable assignment.

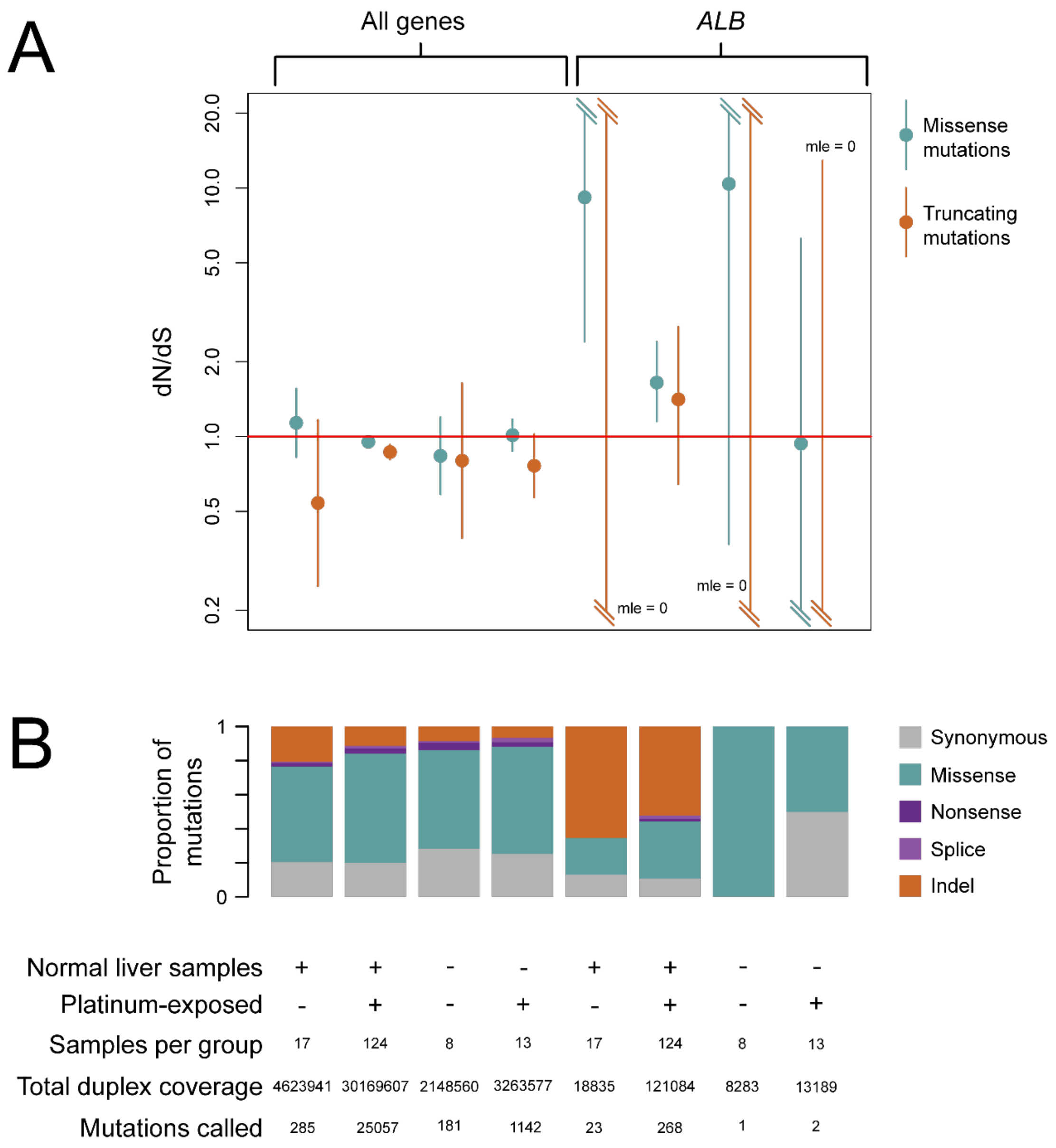


**Fig. S10. dNdS ratio of albumin gene.**

**(A)** dN/dS ratios for missense and truncating mutations across the whole genome and over just the gene encoding albumin (*ALB*) in liver and non-liver samples with and without platinum exposure. The central point represents the maximum likelihood estimate (MLE), and the bars represent the 95% confidence interval.

**(B)** The proportion of mutations in each sample group that are missense, nonsense, synonymous, truncating, or splice site mutations. The sample group is shown below, with the number of samples included in each group, the total duplex coverage (the mean per-gene coverage summed across all samples within the sample group, summed across all genes in the 'all genes' analysis and just for *ALB* in the albumin-only analysis), and the total number of mutations that could be assessed by the dndscv algorithm.
